# The interplay between oxidative stress and inflammation supports autistic-related behaviors in mice

**DOI:** 10.1101/2023.09.11.557109

**Authors:** Luca Pangrazzi, Enrica Cerilli, Luigi Balasco, Caterina Tobia, Ginevra Matilde Dall‘O’, Gabriele Chelini, Samuel Perini, Michele Filosi, Teresa Ravizza, Annamaria Vezzani, Giovanni Provenzano, Anna Pastore, Birgit Weinberger, Enrico Domenici, Yuri Bozzi

## Abstract

Autism Spectrum Disorder (ASD) is a highly prevalent neurodevelopmental condition characterized by social communication deficits and repetitive/restricted behaviors. Several studies showed that inflammation may contribute to ASD. Here we used RT-qPCR, RNA sequencing, immunohistochemistry, and flow cytometry to show that pro-inflammatory molecules were increased in the cerebellum and periphery of mice lacking *Cntnap2* (*Cntnap2^−/−^*), a robust model of ASD. In parallel, oxidative stress was present in the cerebellum of mutant animals. Systemic treatment with N-acetyl-cysteine (NAC) rescued cerebellar oxidative stress and inflammation as well as motor and social impairments in *Cntnap2^−/−^* mice. This was accompanied by improved function of microglia cells in NAC-treated mutant animals. Intriguingly, social deficits, cerebellar inflammation and microglia dysfunction were induced by NAC in *Cntnap2^+/+^*animals. Our findings therefore suggest that the interplay between oxidative stress and inflammation may support ASD-related behaviors in mice.

## Introduction

Autism Spectrum Disorder (ASD) represents a heterogeneous group of neurodevelopmental syndromes characterized by deficits in social communication and social interaction, as well as repetitive patterns of behavior, interests or activities (1). Multiple comorbidities, including attention-deficit/hyperactivity disorder (ADHD), depression and anxiety, obsessive-compulsive disorder (OCD), epilepsy and immune and autoimmune dysregulation, can be typically found in ASD. Worldwide, it has been estimated that about 1/100 children are diagnosed with ASD (2). Many studies have focused on the identification of genes associated with ASD and in proposing strategies of interventions. Despite this, considerable work is still needed to identify effective medical treatments.

Pro-inflammatory dysfunction and impairments in immune cells are known to be associated with ASD (3). Studies performed in humans showed that higher levels of pro-inflammatory molecules interleukin (IL)-1β, IL-6, IL-8, IL-12p40, tumour necrosis factor (TNF), and IL-17 were present in the plasma of medication-free ASD children, in comparison with age-matched healthy controls (4,5). In addition, the expression of cytokines interferon (IFN)γ, IL-1β, IL-6, TNF and chemokine C-C motif ligand (CCL)-2) was increased in the brains and cerebrospinal fluid of ASD subjects (6,7). Similarly, elevated levels of pro-inflammatory molecules have been described in the brain and in the peripheral blood of mouse models of ASD (2).

Alongside with inflammation, the accumulation of oxygen radicals, which causes the onset of oxidative stress, has been proposed to support ASD severity and pathogenesis. Several reports have linked ASD to increased levels of ROS and decreased antioxidant capacity (8–10). In parallel, parameters related to oxidative stress measured in the periphery have been proposed as biomarkers for ASD in humans (11,12). A meta-analysis we recently performed using the dbMDEGA database, which includes transcriptomic data from 14 mouse models of ASD, revealed that the expression of genes coding for enzymes involved in the ROS scavenging system was reduced in the brain of ASD mice (3). Since strong connections between oxidative stress and inflammation have been widely documented (13,14), evidence exists that the interplay inflammation-oxidative stress may be a key contributor to ASD-related behaviors.

Contactin Associated Protein 2 knockout (*Cntnap2^−/−^*) mice have been proposed an optimal mouse model to study ASD, since deficits in the two core ASD behavioral domains are present in these animals (15). In humans, mutations in CNTNAP2 are associated with a neuropsychiatric and neurological syndrome, characterized by ASD, epilepsy, and speech and language disorder ASD (16).

In the current study, we investigated whether pro-inflammatory changes and oxidative stress may be present in the brain of *Cntnap2^−/−^* mice. Increased levels of ROS and inflammation were identified in the cerebellum of mutant mice, as well as in the periphery. Administration of the antioxidant molecule N-acetyl-cysteine (NAC), rescued motor and social behaviours in mutant mice, along with inflammation in the cerebellum and the periphery. Intriguingly, NAC treatment induced social deficits, oxidative stress, and pro-inflammatory dysfunction in the cerebellum of *Cntnap2^+/+^* animals.

Taken together, our work described that an interplay between oxidative stress and inflammation in the cerebellum may support ASD-related behaviors in mice.

## Results

### 1. Pro-inflammatory molecules increase in the cerebellum of *Cntnap2^−/−^* mice

We first analysed the expression of pro-inflammatory molecules IL-6, TNF, IFNγ, and IL-1β in the cerebral cortex, cerebellum and hippocampus of mutant and *Cntnap2^+/+^*adult (6-9 months) mice (Fig. 1a-d). Overall, mRNA expression of all cytokines was higher in the cerebellum, in comparison with the other brain regions. Furthermore IL-6, TNF, IFNγ and IL-1β levels increased in the cerebellum of *Cntnap2^−/−^* mice, in comparison with *Cntnap2^+/+^*animals. To assess the possible contribution of immune cells to pro-inflammatory conditions, we measured the expression of IL-6, TNF, and IFNγ in immune cells using flow cytometry (Fig. 1e-g). Gating strategy is shown in Fig. S1. Higher levels of IL-6 and TNF were present in CD45^+^ cells within the cerebellum, and a significant increase was found in *Cntnap2^−/−^* mice compared to *Cntnap2^+/+^*controls (Fig. 1e,f). IFNγ is mostly expressed by T cells, which infiltrate the brain and are maintained there as tissue-resident cells (17). Again, IFNγ levels were significantly higher in the *Cntnap2^−/−^* cerebellum (Fig. 1g.) Similarly IL-2, a T cell survival and proliferation factor produced by activated T cells, was more expressed in the cerebellum of *Cntnap2^−/−^* mice (Fig. 1h). mRNA levels of CCL3 and CCL8 chemokines (which regulate the migration of immune cells to site of inflammation; 18), and metalloproteases MMP3 and MMP12 (which are associated to tissue damage and response to stress and thus directly connected with inflammation) were significantly overexpressed in the cerebellum but not in other areas of *Cntnap2^−/−^* mice (Fig. 1i-l). Other molecules related to inflammation (IFNβ, TLR1, TLR2, TLR5, TLR6, TLR9, CCL8, CCL21A, CCL20) and tissue damage (MMP8, MMP13) showed a similar trend, being more expressed in the cerebellum of mutant animals compared to *Cntnap2^+/+^* controls (Table S1). Cerebellar expression of IL-15, TLR3, TLR4, and CCL2 mRNAs did not differ between genotypes. Other areas relevant to ASD, such as the prefrontal cortex and striatum, showed low and comparable mRNA levels of proinflammatory cytokines and metalloprotease in both genotypes (data not shown). Taken together, our results clearly described that increased levels of molecules related to inflammation and supporting tissue damage were only present in the cerebellum of *Cntnap2^−/−^* mice.

**Figure 1.**
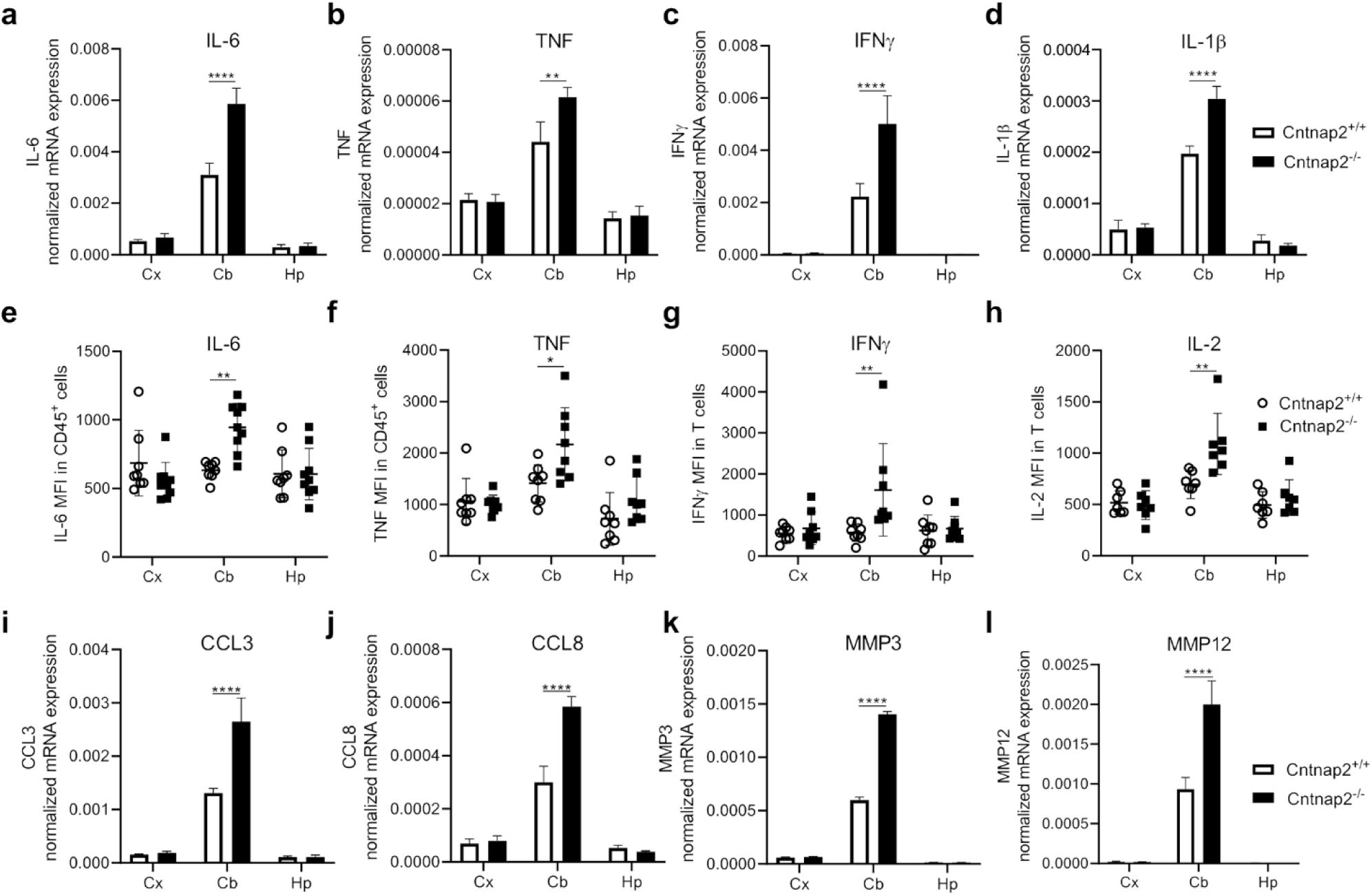
Pro-inflammatory changes in the brain of *Cntnap2^+/+^* and *Cntnap2^−/−^* mice. mRNA expression of (**a**) IL-6, (**b**) TNF, (**c**) IFNγ, and (**d**) IL-1β in the cerebral cortex (Cx), cerebellum (Cb) and hippocampus (Hp) assessed using RT-qPCR. Mean fluorescence intensity (MFI) of (**e**) IL-6, (**f**) TNF, (**g**) IFNγ, and (**h**) IL-2 in the Cx, Cb, and Hp of *Cntnap2^+/+^* and *Cntnap2^−/−^* mice measured using flow cytometry. mRNA expression of (**i**) CCL3, (**j**) CCL8, (**k**) MMP3 and (**l**) MMP12 in the Cx, Cb and Hp assessed using RT-qPCR. Two-way ANOVA, Tukey post-hoc test. n a-d and n i-l = 8; n e-h = 7-10 in each group. *p<0.05; **p<0.01; ***p<0.001; ****p<0.0001.

### 2. Large-scale transcriptomic analysis highlights pro-inflammatory changes and oxidative stress in the cerebellum of *Cntnap2^−/−^* mice

To study large-scale mRNA expression in the cerebellum of *Cntnap2^−/−^*mice, we performed a RNA sequencing (RNAseq) analysis. To identify significant differentially expressed genes, an adjusted p value (p_adj_) < 0.05 and a fold change (FC) > 0.35 or < 0.35 were used as cut-off. In this way, 589 and 363 genes were upregulated or downregulated respectively in *Cntnap2^−/−^* mice, in comparison to *Cntnap2^+/+^* animals. Most over-expressed transcripts coded for calcium binding proteins, ribosomal proteins as well as molecules involved in inflammatory responses (Table S2). Most downregulated transcripts were involved in the regulation of neural functions (Table S3). Importantly, while genes involved in inflammation (*i.e.* Ccr1, Pla2g4c, Il22, Il22b, Ccl12, Ctsw, and Ccl27a) were upregulated, antioxidant enzymes Cat and Sod3 were downregulated in *Cntnap2^−/−^*animals (Table S4). Interestingly, complement genes C4b, C1qtnf1 and C3 were less expressed by mutant mice, in comparison to *Cntnap2^+/+^* controls (Table S4).

As a next step, gene set enrichment analysis (GSEA) was performed on significant up- and down-regulated genes (Fig. 2b-c, Fig. S2a,b). Top significantly enriched gene ontology (GO) were represented by ribosome and mitochondrial components, including “structural constituent of ribosome”, “ribosome”, “mitochondrial ribosome” as well as “mitochondrial inner membrane” (Fig.2b, Fig.S2a). Importantly, all ontologies were activated in the cerebellum of mutant mice. Top KEGG pathways included “ribosome”, “oxidative phosphorylation”, pathways involved in immune responses (COVID-19, Herpes simplex virus 1 infection, Human papillomavirus infection), and neural functions (calcium signaling pathways and axon guidance) (Fig. 2c, Fig.S2b). While most immune-related pathways and pathways related with oxidative stress were positively enriched, neural pathways were suppressed in mutant animals (Fig. 2b,c, Fig.S2a,b).

**Figure 2.**
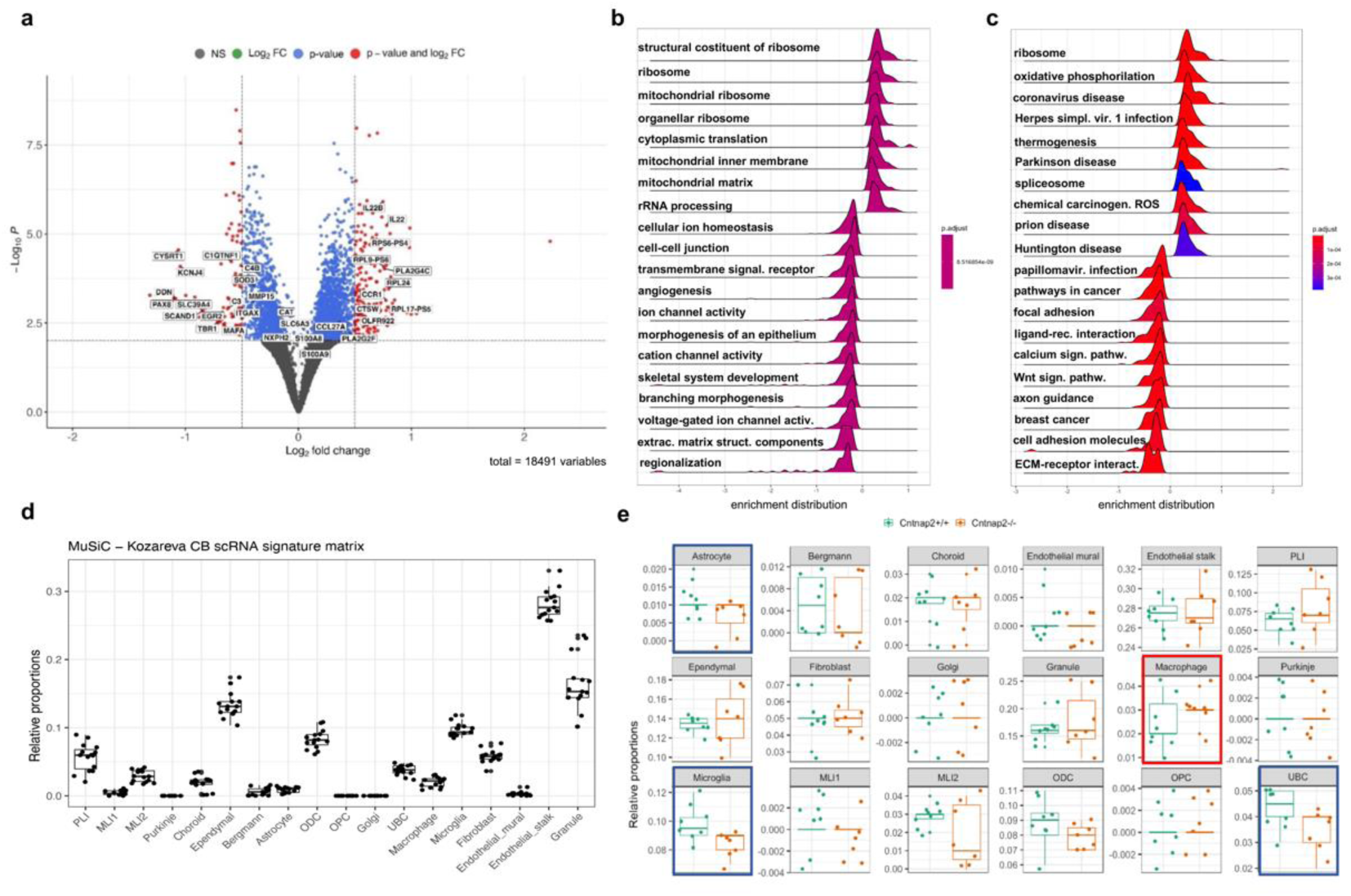
RNAseq analysis in the cerebellum of Cntnap2^+/+^ and Cntnap2^−/−^ mice. (**a**) Volcano plot showing some of the most differentially expressed genes. Most represented gene ontologies (GO) (**b**) and KEGG pathways (**c**) enriched in Cntnap2^−/−^ mice. P values are reported in the graphs using the scale color. (**d**) Cell proportions within the cerebellum inferred by the MuSiC algorithm using the signature matrix of Kozareva *et al.* (19) as a reference. (**e**) Relative cell proportions of cerebellar cell types within *Cntnap2^+/+^* and *Cntnap2^−/−^* mice. Kruskal-Wallis test, n=8 in each group. Red square p<0.05, blue squares p<0.01.

To track compositional alterations of cell types, a cellular deconvolution approach was performed on our RNAseq dataset (Fig. 2d-e, Fig. S3a,b). Cell proportions within the cerebellum were inferred by the MuSiC algorithm, using the signature matrix of Kozareva *et al.* as a reference (19). Relative proportions of cerebellar cell types are shown in Fig. 2d. Most represented cell types were endothelial cells, granule, ependymal cells and microglia. When relative proportions were compared, unipolar brush cells (UBC), astrocytes, and microglia cells were reduced, while marcophages were increased in *Cntnap2^−/−^* mice, in comparison to *Cntnap2^+/+^* controls (Fig. 2e). We next assessed the contribution of the variation in proportions of astrocytes, macrophages, and microglia on the difference in gene expression identified between the two genotypes. We therefore performed GO and KEGG pathways analysis after adjusting for each subset (astrocytes, macrophages, and microglia) independently, the macrophages/microglia (Macro-Micro) group and the astrocyte/macrophage/microglia (Astro-Macro-Micro) group (Fig. S3a,b). Overall, components enriched in the GO analysis were similar when the groups Macro-Micro, macrophages, and microglia were compared, while they were different from the astrocyte group (Fig. S3a,b). No GO terms were significantly enriched in the astrocyte group. When KEGG pathways were assessed, two pathways related to inflammation (“Th17 differentiation” and “Inflammatory bowel disease”) were enriched after correcting for the Astro-Macro-Micro group. Furthermore, “Ribosome” as well as pro-inflammatory pathways (“Inflammatory bowel disease” and “COVID-19”) were enriched in the Macro-Micro, Macrophage and Microglia groups, while no pathways were identified for the astrocyte group alone. In summary, our analysis indicates that pro-inflammatory processes are induced in the cerebellum of *Cntnap2^−/−^* mice, while the cellular responses to oxidative stress may be compromised.

### 3. Pro-inflammatory changes are present in the PB of *Cntnap2^−/−^* mice

Increased expression of pro-inflammatory molecules was described in the peripheral blood (PB) of patients with ASD (3–5). We therefore investigated whether systemic pro-inflammatory changes may be present not only in the cerebellum, but also in the PB of *Cntnap2^−/−^* mice. We assessed the levels of IL-6, TNF and IFNγ in the whole PB mononuclear cells (PBMC) population, monocytes and T cells obtained from adult *Cntnap2^−/−^* and *Cntnap2^+/+^* animals (Fig. 3). Gating strategy used to define the populations of interest in flow cytometry experiments is reported in Fig. S1b. Frequency of CD14^+^ monocytes was comparable in mutant and control mice, while T cells were increased in *Cntnap2^−/−^*animals (Fig. 3a,b). In parallel, CD8^+^ T cells increased while CD4^+^ T cells were decreased in mutant mice (Fig. 3c,d). No differences between the groups were observed in the frequencies of CD44^−^ CD62L^+^ naïve, CD44^+^CD62L^+^ central-memory and CD44^+^CD62L^−^ effector-memory CD8^+^ and CD4^+^ T cells (data not shown). The expression of IL-6 was increased in PBMCs and monocytes from *Cntnap2^−/−^* mice, in comparison with the same populations in control animals, while it did not change in T cells (Fig. 3e-g). Again, TNF was overexpressed by mutant animals within monocytes, but it did not change in T cells and in the whole PBMC population (Fig. 3h-j). In addition, the expression of IFNγ in *Cntnap2^−/−^* mice was higher in PBMCs, as well as in T cells, when compared with the same cell populations in *Cntnap2^+/+^*mice (Fig. 3k-n). No differences were found for IFNγ expression in monocytes (Fig. 3o). Thus, our results demonstrated that pro-inflammatory changes in *Cntnap2^−/−^*mice are present not only in the cerebellum, but also in the PB.

**Figure 3.**
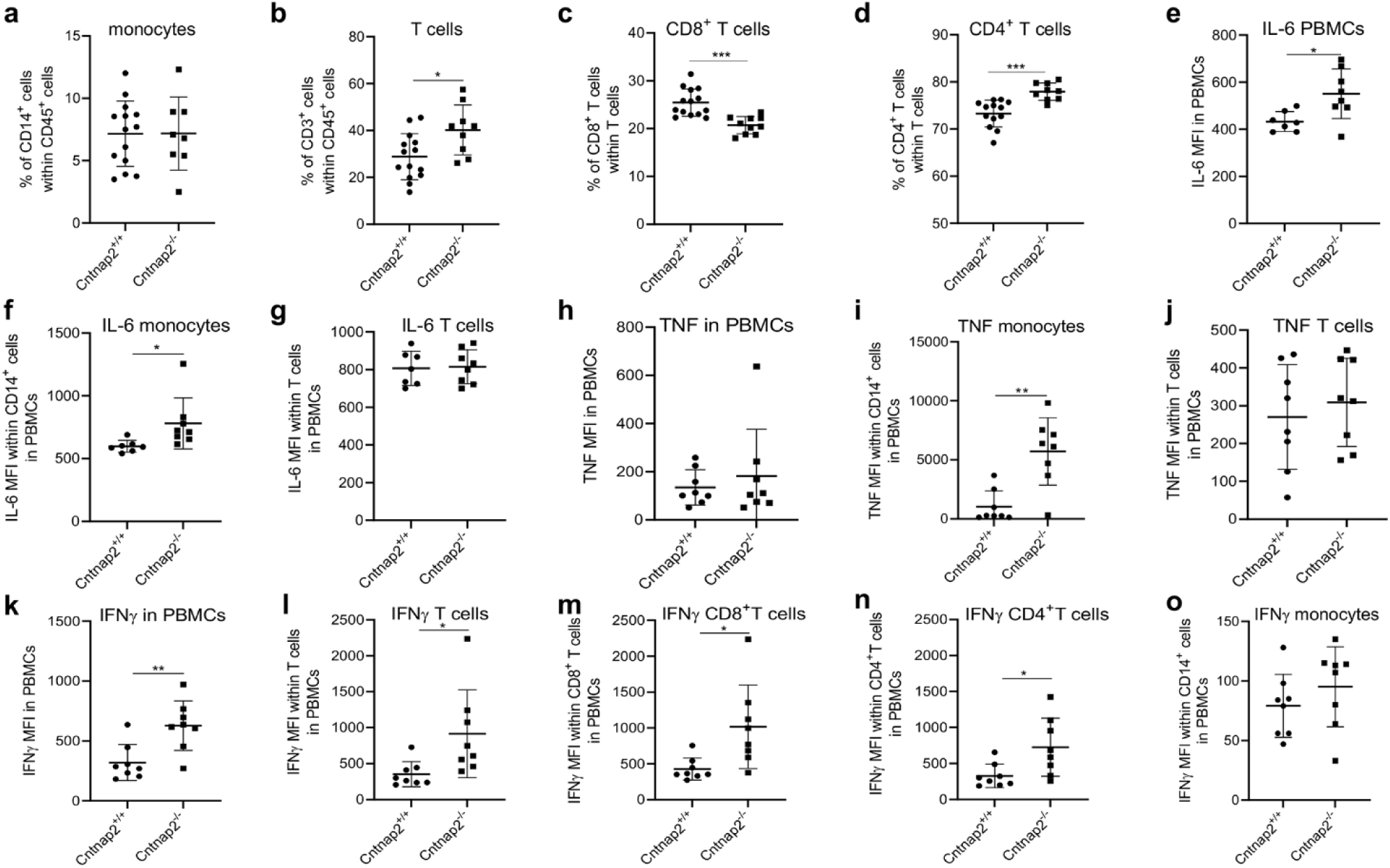
Pro-inflammatory changes in the peripheral blood of *Cntnap2^+/+^* and *Cntnap2^−/−^* mice. (**a**) Frequency of CD14^+^ cells (monocytes) and (**b**) T cells within CD45^+^ immune cells measured using flow cytometry. Frequency of (**c**) CD8^+^ and (**d**) CD4^+^ cells within T cells. n_Cntnap2+/+_=14, n_Cntnap2+/+_=9. Mean fluorescence intensity (MFI) of IL-6 in (e) PBMCs, (**f**) monocytes and (**g**) T cells. n_Cntnap2+/+_=7, n_Cntnap2+/+_=8. MFI of TNF in (**h**) PBMCs, (**i**) monocytes and (**j**) T cells. MFI of IFNγ within (**k**) T cells, (**m**) CD8^+^ T cells, (**n**) CD4^+^ T cells and (**o**) monocytes. N c_ntnap2+/+_=8, n_Cntnap2+/+_= 8. Unpaired t test. *p<0.05; **p<0.01; ***p<0.001.

### 4. N-acetyl-cysteine improves ASD-related behaviors in *Cntnap2^−/−^* mice but has detrimental effects in *Cntnap2^+/+^* animals

To understand whether a link may exist between oxidative stress, inflammation and ASD-related behaviors in *Cntnap2^−/−^* mice, we aimed at targeting cerebellar oxidative stress and inflammation *in vivo*. As N-acetyl-cysteine (NAC) shows both antioxidant and anti-inflammatory properties and its capability to cross blood-brain barrier has been documented (20), we treated *Cntnap2^−/−^* and *Cntnap2^+/+^* mice systemically with NAC or vehicle (PBS) for 28 days. Motor (open field and rotarod test) and social (3-chamber social test) behaviors were tested on NAC- and PBS-treated *Cntnap2^−/−^*and *Cntnap2^+/+^* mice (Fig. 4). As already described (15), PBS-treated *Cntnap2^−/−^*mice were hyperactive, showing increased mean velocity and distance travelled in the open field test in comparison to *Cntnap2^+/+^* animals (Fig. 4a). Interestingly, mutant animals treated with NAC displayed a rescue of both parameters, which were then similar to *Cntnap2^+/+^* PBS-treated mice. No differences were observed between PBS- and NAC-treated *Cntnap2^+/+^* mice. Similarly, PBS-treated *Cntnap2^−/−^*mice spent significantly more time on the rotarod and concluded the test with higher RPM of the rotarod machine when compared with PBS-treated *Cntnap2^+/+^*mice (Fig. 4b). Again, NAC-treated mutant animals performed similarly to *Cntnap2^+/+^* mice. In order to assess the effects of NAC on social behavior, sociability of Cntnap2^+/+^ and *Cntnap2^−/−^* mice was assessed in the 3-chamber test (Fig. 4C). All experimental groups spent a similar time in the center chamber (Fig. S4). When compared with PBS-treated *Cntnap2^+/+^* animals, PBS-treated *Cntnap2^−/−^*mice showed reduced preference for the mouse chamber (Fig. 4c and Fig. S4). Conversely, NAC-treated *Cntnap2^−/−^* mice showed increased sociability, as their sociability index was comparable to that of PBS-treated *Cntnap2^+/+^* mice (Fig. 4c and Fig. S4). Unexpectedly, NAC-treated *Cntnap2^+/+^* mice showed reduced preference for the mouse chamber compared to the PBS-treated Cntnap2^+/+^ group (Fig. 4c and Fig. S4).

**Figure 4.**
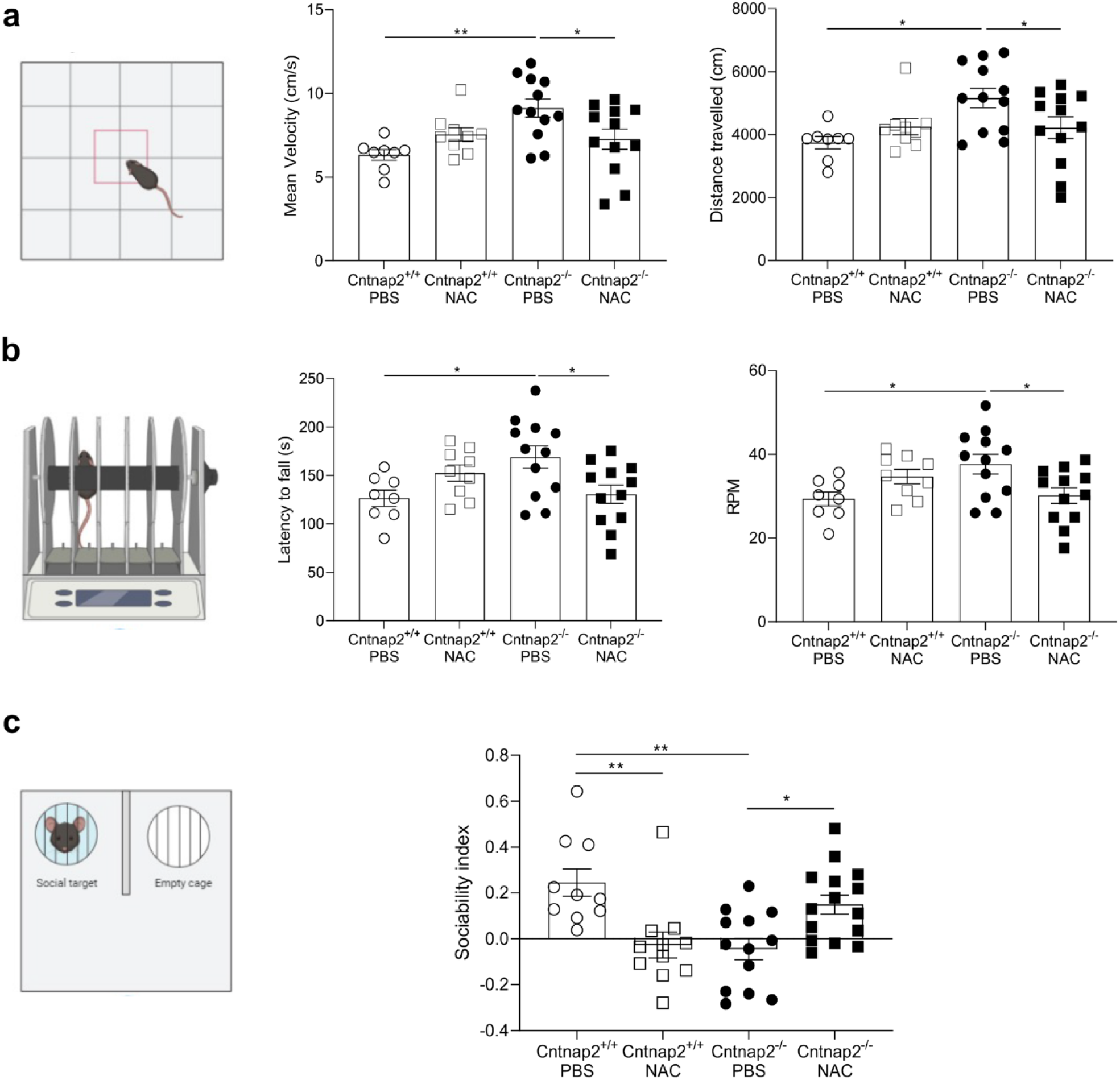
Behavioral tests in *Cntnap2^+/+^* and *Cntnap2^−/−^* mice treated with NAC. (**a**) Mean velocity (cm/s) and distance travelled (cm) in the open field test in NAC-treated *Cntnap2^+/+^* and *Cntnap2^−/−^* mice and PBS-treated control animals of both genotypes. (**b**) Latency to fall (s) and revolutions per minute (RPM) in the rotarod test. (**c**) Sociability index (time in mouse chamber/time in mouse+empty chambers) in the 3-chamber test. n_Cntnap2+/+PBS=_8, n_Cntnap2+/+NAC=_9, n_Cntnap2−/−PBS=_12, n_Cntnap2+/+NAC=_12. Two-way ANOVA, Tukey post-hoc test. *p<0.05; **p<0.01.

Taken together, our results clearly showed that NAC treatment improved both motor and social behaviours of *Cntnap2^−/−^* mice, while NAC induced social impairments in *Cntnap2^+/+^* animals.

### 5. NAC reduces oxidative stress and inflammation in the cerebellum of *Cntnap2^−/−^* mice and increases ROS and pro-inflammatory molecules in *Cntnap2^+/+^* animals

We next assessed whether behavioral changes induced by NAC may be accompanied by differences in the expression of molecules related to oxidative stress and inflammation in the cerebellum (Fig. 5). Levels of reactive oxygen species (ROS) were higher in *Cntnap2^−/−^* mice treated with PBS compared with PBS-treated *Cntnap2^+/+^* animals (Fig. 5a). After NAC treatment, ROS levels were significantly decreased in mutant mice, while no differences were found between NAC- and PBS-treated *Cntnap2^+/+^* animals. When ROS were measured within cerebellar neurons, similar results were observed for *Cntnap2^−/−^*mice (Fig. 5b). Despite this, NAC-treated *Cntnap2^+/+^*mice showed higher ROS levels in comparison with PBS-treated Cntnap2^+/+^ animals (Fig. 5b). No differences between groups were found when ROS were measured within microglia and T cells (data not shown). We then measured the levels of molecules related to inflammation and tissue damage within the cerebellum in the four experimental groups (Fig. 5c-i and Table S5). In agreement with what previously observed in untreated mice (Fig. 1a), IL-6 mRNA was higher in PBS-treated *Cntnap2^−/−^* mice compared with PBS-treated *Cntnap2^+/+^* animals (Fig. 5c). After NAC treatment, IL-6 mRNA expression in mutant mice was significantly decreased and comparable with IL-6 levels measured in *Cntnap2^+/+^* mice (Fig. 5c). In agreement with the toxic effects of NAC already observed in *Cntnap2^+/+^* animals at behavioral level (Fig. 4), increased IL-6 mRNA expression was observed in NAC-treated *Cntnap2^+/+^* mice compared with PBS-treated *Cntnap2^+/+^* littermates. Similar results were found when mRNA levels of IL-1β, TNF, IFNγ, CCL3, MMP3, IL-22, and other molecules related to inflammation and tissue damage were measured in the four groups (Fig. 5d-i and Table S5). In parallel, most of the tested mRNAs increased their expression in the cerebellum when *Cntnap2^+/+^* mice were treated with NAC (Fig. 5 and Table S5). We finally assessed the expression of IL-6, TNF and IFNγ in immune cell subsets within the cerebellum, *i.e* the CD14^+^ cell subset including activated microglia and macrophages, and T cells (Fig. 5j-l). IL-6 expression measured in CD14^+^ immune cells was higher in mutant animals, and decreased after NAC treatment (Fig. 5j). Similar results were observed for TNF expression in CD14^+^ cells and IFNγ in T cells (Fig. 5k,l). Again, an overexpression of IL-6 was observed in *Cntnap2^+/+^*mice treated with NAC compared with PBS-treated *Cntnap2^+/+^* controls (Fig. 5j).

**Figure 5.**
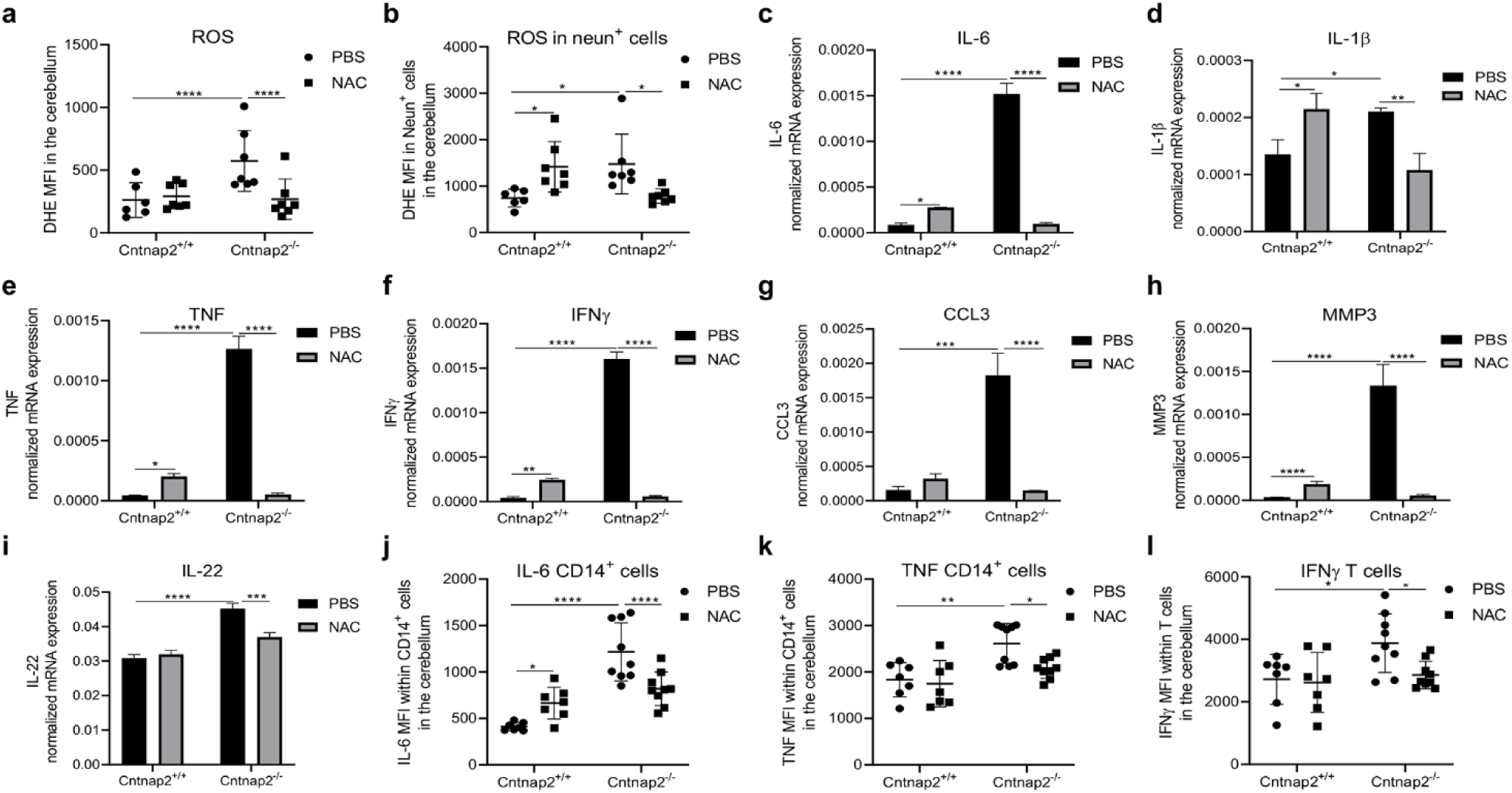
ROS levels and pro-inflammatory molecules in the cerebellum of *Cntnap2^+/+^* and *Cntnap2^−/−^* mice treated with NAC. ROS levels (=DHE MFI) in (**a**) the cerebellum and (**b**) cerebellar neurons (Neun^+^ cells) measured using flow cytometry. n_Cntnap2+/+PBS=_6, n_Cntnap2+/+NAC=_7, n_Cntnap2−/−PBS=_7, n_Cntnap2+/+NAC=_7. mRNA expression of (**c**) IL-6, (**d**) IL-1β, (**e**) TNF, (**f**) IFNγ, (**g**) CCL3, (**h**) MMP3, and (**i**) IL-22 assessed using RT-qPCR. mRNA expression was normalized against the housekeeping gene β-actin. n=8 in each group. MFI of (**j**) IL-6 in CD14^+^ cells, (**k**) TNF in CD14^+^ cells, and (**l**) IFNγ, in T cells measured using flow cytometry. n=7-9 in each group. Two-way ANOVA, Tukey post-hoc test. *p<0.05; **p<0.01; ***p<0.001, ****p<0.0001.

Taken together, our results indicate that oxidative stress and inflammation in the cerebellum of *Cntnap2^−/−^* mice are strongly counteracted by NAC treatment. Conversely, NAC treatment in *Cntnap2^+/+^* mice results in increased levels of oxygen radicals and molecules related to inflammation and tissue damage.

### 6. NAC improves microglia function in *Cntnap2^−/−^* mice while impairing it in *Cntnap2^+/+^*controls

As microglia is a key regulator of pro-inflammatory processes, and changes related to inflammation and oxidative stress were identified within these cells in in the *Cntnap2^−/−^* cerebellum mice (Fig. 2d,e and Fig. S3), we assessed the phenotype of microglia cells in the cerebellum of NAC- and PBS-treated *Cntnap2^−/−^* and *Cntnap2^+/+^* mice (Figure 6). Crus 1 and Crus 2 cerebellar areas (Fig. S5a) have been implicated in social behaviors in both humans (21) and mouse models (22; 23). We therefore assessed the frequency and the branching of Iba-1^+^ / Tmem119^+^ microglia cells in the molecular (ML) and granular (GL) layers of Crus 1 and Crus 2 from PBS- and NAC-treated *Cntnap2^−/−^* and *Cntnap2^+/+^* mice (Fig. 6 and Fig. S5a,b). In line with our RNAseq results (Fig. 2d,e and Fig. S3), the frequency of microglia cells was decreased in PBS-treated mutant mice compared to PBS-treated controls in Crus 2 (Fig. 6a,b) and Crus 1 (Fig. S5c). Microglia cell counts Crus1 (Fig. 5Sc) and Crus 2 (Fig. 6b) decreased in NAC-treated *Cntnap2^+/+^* animals compared to PBS-treated *Cntnap2^+/+^* mice, again indicating a potentially toxic effect of NAC in *Cntnap2^+/+^* animals. Conversely, no differences were observed in mutant mice after NAC administration. We next performed a Sholl analysis to investigate the branching of microglia cells in Crus 1 and Crus 2 areas. Significant decrease in the branching of microglia cells was observed in PBS-treated *Cntnap2^−/−^* mice and in NAC-treated *Cntnap2^+/+^*animals, compared to PBS-treated *Cntnap2^+/+^* controls (Fig. S5c and Fig. 6c). Interestingly, microglia branching in mutant mice was completely rescued by NAC treatment in Crus 1 and Crus 2 (Fig. S5c and Fig.6c).

**Figure 6.**
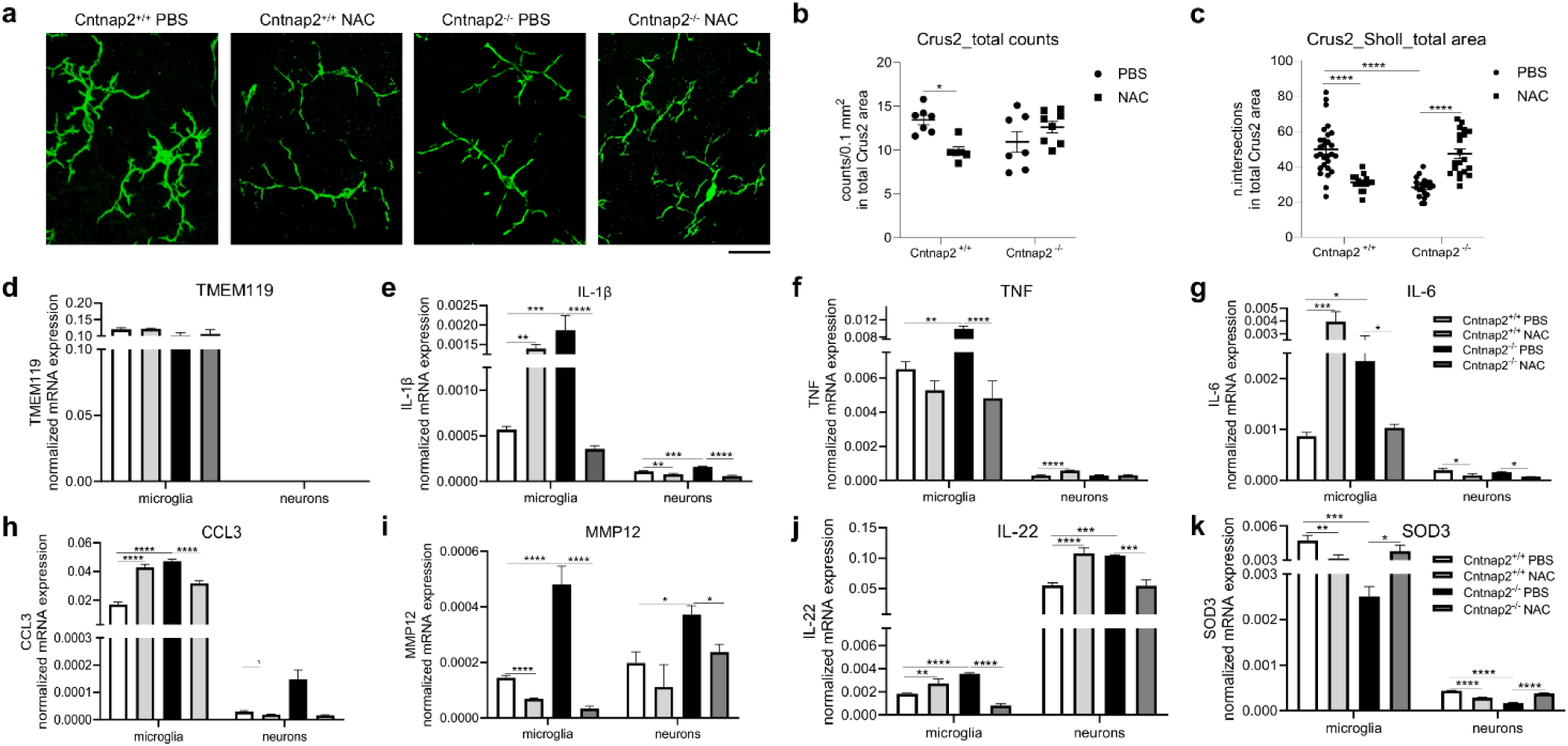
The impact of NAC on microglia function in the cerebellum of *Cntnap2^+/+^* and *Cntnap2^−/−^* mice. **(a)** Representative pictures showing microglia (iba-1^+^) cells in the Crus2 area of PBS- and NAC-treated *Cntnap2^+/+^* and *Cntnap2^−/−^* mice. Scalebar = 500µM. Numbers of (**b**) iba-1^+^ cells (expressed in 0.1 mm^2^) and (**c**) intersections calculated with the Sholl analysis in the Crus2 total area of PBS- and NAC-treated animals. For cell number quantification: n_Cntnap2+/+PBS=_7, n_Cntnap2+/+NAC=_6, n_Cntnap2−/−PBS=_7, n_Cntnap2+/+NAC=_8; for Sholl analysis: n_Cntnap2+/+PBS=_14, n_Cntnap2+/+NAC=_7, n_Cntnap2−/−PBS=_9, n_Cntnap2+/+NAC=_10. Two-way ANOVA, Tukey post-hoc test. *p<0.05; ****p<0.0001. mRNA expression of (**d**) TMEM119, (**e**) IL-1β, (**f**) TNF, (**g**) IL-6, (**h**) CCL3, (**i**) MMP12, (**j**) IL-22, and (**k**) SOD3 in FACS-sorted microglia cells and neurons from the cerebellum of PBS- and NAC-treated *Cntnap2^+/+^* and *Cntnap2^−/−^* mice assessed using RT-qPCR. n=8 in each group. Two-way ANOVA, Tukey post-hoc test. *p<0.05; **p<0.01; ***p<0.001;****p<0.0001.

To assess whether microglia cells may contribute to pro-inflammatory changes and oxidative stress, Tmem119^+^ microglia cells and, as a control, Neun^+^ neurons were FACS-sorted from the cerebellum of NAC- and PBS-treated *Cntnap2^+/+^* and *Cntnap2^−/−^* mice. mRNA expression of molecules related to inflammation and oxidative stress was assessed in sorted microglia cells and neurons (Fig. 6d-k). Importantly, all Iba-1^+^ cells in the cerebellum were Tmem119^+^ (Fig. S5b). The purity of the Tmem119^+^ fraction was confirmed (Fig. 6d). Overall, levels of pro-inflammatory molecules and chemokines were higher in microglia cells in comparison to neurons, with the exception of IL-22 (Fig. 6j). In the Tmem119^+^ fraction, PBS-treated *Cntnap2^−/−^* mice showed increased levels of IL-1β, TNF, IL-6, CCL3, and MMP12 in comparison to PBS-treated *Cntnap2^+/+^*animals, and they were all reduced after the NAC administration (Fig. 6e-h). In parallel, some of these molecules (IL-1β, IL-6 and CCL3) were increased in NAC-treated *Cntnap2^+/+^* mice (Fig. 6e,g,h). Similar results were observed in neurons, although only TNF was upregulated in this fraction in NAC-treated *Cntnap2^+/+^*mice (Fig. 6f). While the expression of MMP12 was similar between microglia and neurons, IL-22 was more expressed by Neun^+^ cells (Figure 6I,j). Despite this, IL-22 expression in the four experimental groups was similar to that of the other cytokines (Fig. 6e,f,g,j). Finally, SOD3 mRNA expression (that was low in PBS-treated mutant animals), was induced in *Cntnap2^−/−^*microglia and neurons after NAC treatment, while decreasing in NAC-treated *Cntnap2^+/+^*mice (Fig.6k). Taken together, these results show that NAC improves the phenotype of microglia cells and counteract pro-inflammatory dysfunction in the cerebellum of *Cntnap2^−/−^* mice, while inducing microglia impairment, inflammation, and oxidative stress in *Cntnap2^+/+^*animals.

### 7. NAC reduces ROS and pro-inflammatory cytokines in the peripheral blood, bone marrow, and spleen of *Cntnap2^+/+^* and Cntnap2^−/−^ mice

We then investigated whether NAC treatment may additionally affect the levels of oxygen radicals and pro-inflammatory molecules in the periphery (Fig. 7). Similar to what observed in the cerebellum, ROS levels were higher in the blood from PBS-treated *Cntnap2^−/−^* mice compared to PBS-treated *Cntnap2^+/+^* animals (Fig. 7a). After NAC treatment, ROS were reduced in mutant animals while they increased in *Cntnap2^+/+^*mice (Fig. 7a). We next measured reduced glutathione (GSH), cysteine (cys), and cysteinyl-glycine (cys-gly) in the serum. While GSH and cys levels were similar in the four experimental groups, cys-gly was increased in *Cntnap2^+/+^* mice after NAC treatment. No significant differences were observed between PBS- and NAC-treated mutant animals (Fig. 7b-d). In addition, we assessed the expression of IL-6 and TNF in monocytes and T cells, as well as IFNγ in T cells (Fig. 7e-i). Within monocytes, IL-6 and TNF levels were higher in PBS-treated mutant mice and were restored to basal levels after NAC treatment (Fig. 7e,f). No differences between the four experimental groups were observed when IL-6 and TNF expression was analysed within T cells (Fig. 7g,h). In addition, the levels of IFNγ within T cells were higher in PBS-treated *Cntnap2^−/−^* mice compared with PBS-treated *Cntnap2^+/+^* animals, and decreased in mutant mice after NAC treatment (Fig. 7i). Similar results were observed for IFNγ levels measured within CD8^+^ and CD4^+^ T cells (Fig. S6a,b).

**Figure 7.**
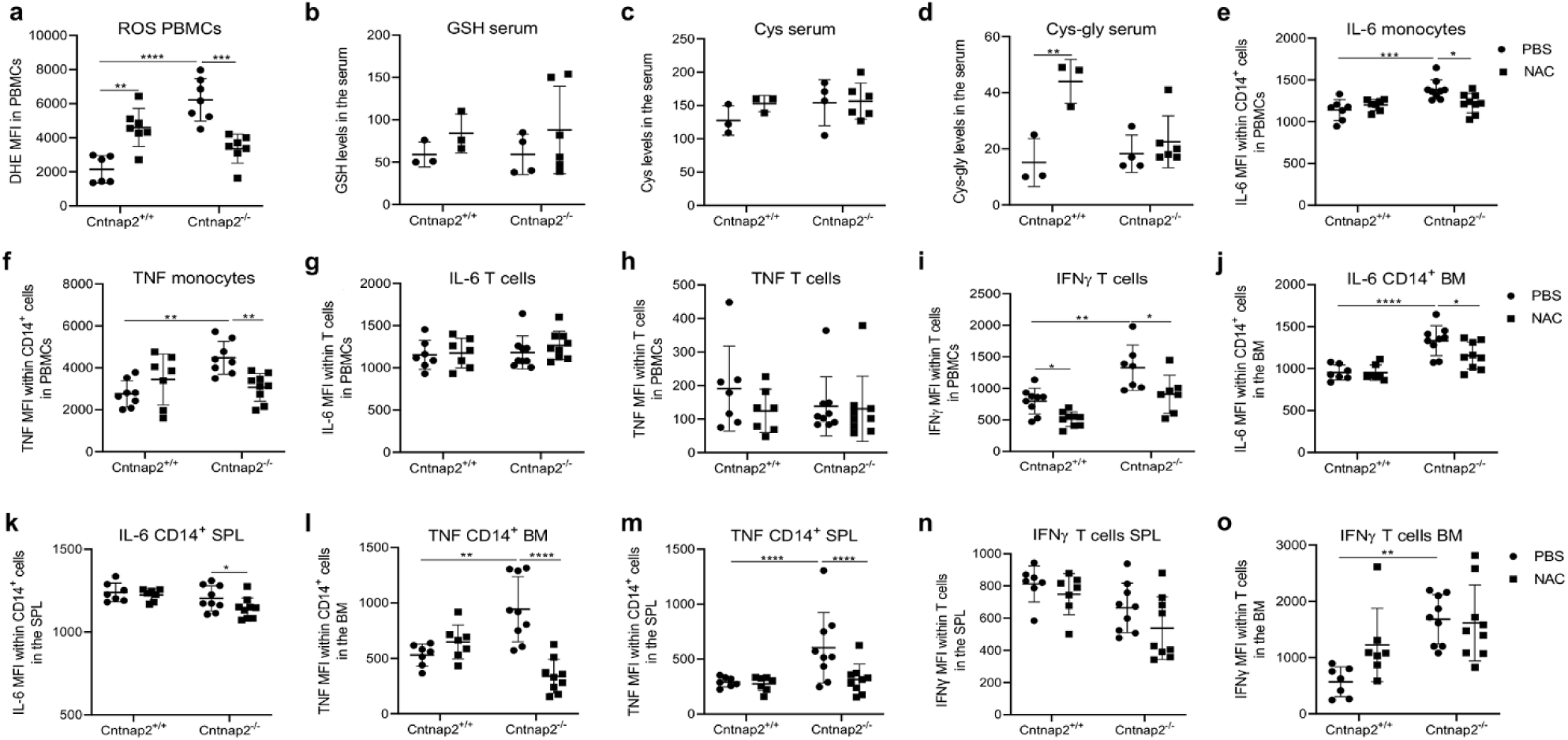
The impact of NAC on ROS and pro-inflammatory cytokines in the peripheral blood, bone marrow, and spleen of *Cntnap2^+/+^* and *Cntnap2^−/−^* mice. (a) ROS levels (=DHE MFI) in PBMCs from PBS- and NAC-treated Cntnap2^+/+^ and Cntnap2^−/−^ mice. (**b**) GSH, (**c**) Cys, and (**d**) Cys-gly levels in the serum. Mean fluorescence intensity (MFI) of (**e**) IL-6 in monocytes, (**f**) TNF in monocytes, (**g**) IL-6 in T cells, (**h**) TNF in T cells, (i) IFNγ in T cells within PBMCs from PBS- and NAC-treated animals. MFI of (**j**) IL-6 in CD14^+^ cells in the bone marrow (BM), (**k**) IL-6 in CD14^+^ cells in the spleen (SPL), (**l**) TNF in CD14^+^ cells in the bone marrow, (**m**) TNF in CD14^+^ cells in the spleen, (**n**) IFNγ in T cells in the spleen, and (**o**) IFNγ in T cells in the bone marrow. n=3-9 in each group. Two-way ANOVA, Tukey post-hoc test. *p<0.05; **p<0.01; ***p<0.001;****p<0.0001.

We next assessed whether the administration of NAC might additionally affect the levels of pro-inflammatory molecules in the bone marrow and spleen of *Cntnap2^−/−^* and *Cntnap2^+/+^* mice. The bone marrow plays a key role in the generation and the maintenance of innate and adaptive immune cells, and the importance of spleen in immune responses has deeply been investigated. We therefore measured the expression of IL-6, TNF, and IFNγ in CD14^+^ cells and T cells within these two immune organs using flow cytometry (Fig.7j-o). Similar to the PB, IL-6 levels in the bone marrow were higher in PBS-treated *Cntnap2^−/−^* mice, when compared with in PBS-treated *Cntnap2^+/+^* controls (Fig. 7j). No differences between the genotypes were found in the spleen (Fig. 7k). In *Cntnap2^−/−^*mice, NAC reduced IL-6 levels in CD14^+^ cells in both organs (Fig. 7j,k) and in T cells from the spleen (Fig. S6d), while IL-6 levels in bone marrow T cells did not differ between the four groups (Fig. S6c). The same results were found for TNF expression in CD14^+^ cells in the bone marrow and spleen (Fig. 7l,m), while TNF levels within T cells did not differ between the four groups (Fig. S6e,f). Finally, IFNγ expression within T cells was higher in the bone marrow of PBS-treated mutant mice, in comparison to PBS-treated *Cntnap2^+/+^*controls. However, NAC treatment did not rescue IFNγ levels in mutant mice, neither in the bone marrow nor in the spleen (Fig. 7n,o).

In summary, pro-inflammatory dysfunction in the PB, bone marrow, and spleen of *Cntnap2^−/−^* mice is counteracted by the administration of NAC. Importantly, pro-inflammatory changes were mostly found within CD14^+^ cells.

## Discussion

ASD is characterized by a broad spectrum of neurodevelopmental conditions affecting children from early childhood and producing clinically significant developmental impairments. Although their exact origin is still unknown, a strong genetic component has been proposed (24). Recent studies documented immune dysfunction, inflammation, and oxidative stress in individuals with ASD and in mouse models (3–13). Thus, despite the high heterogeneity, it is now becoming evident that impairments in the immune system may strongly contribute to the pathogenesis and severity of ASD. In this study, we investigated for the first time the expression of pro-inflammatory molecules in the brain of *Cntnap2^−/−^* mice, a well-established model of ASD. The cerebellum is a brain area important for the motor control of the body. In addition, it plays a fundamental role in some cognitive functions such as emotional control, attention, language, and social behaviors. Several studies have shown that the cerebellum is structurally and functionally abnormal in autistic individuals (25): Individuals with CNTNAP2 mutations present with cerebellar malformations (26), and CNTNAP2 antibodies have been detected in the cerebrospinal fluid of subjects with mild cerebellar ataxia (27, 28). More recently, strong evidence has been obtained supporting the mechanistic contribution of cerebellar dysfunction to the pathogenesis of ASD-like behaviors in mouse models (29). Cerebellar deficits have been described in several mouse models of ASD, including *Cntnap2^−/−^* mice. As an example, Kloth *et al.* reported impaired eyeblink conditioning (a form of associative sensory learning requiring cerebellar plasticity) in *Cntnap2^−/−^* mice (30), which are associated to *Cntnap2* expression pattern in Purkinje and granule cells in the cerebellum (https://mouse.brain-map.org, experiment 544712). In keeping with these findings, a more recent study identified electrophysiological and anatomical alterations in the cerebellum of *Cntnap2^−/−^* mice, at the level of Purkinje cells of the Crus region (31), a key area for the modulation of social behaviors (21–23).

We first showed that in *Cntnap2^−/−^* mice, pathological levels of neuroinflammation were specifically detected within the cerebellum but not within other brain areas relevant to ASD (Fig. 1). We therefore performed an RNAseq study to get an overview about transcriptome differences between *Cntnap2^−/−^* and *Cntnap2^+/+^*mice within the cerebellum. A specific signature of inflammation and oxidative stress was detected in the mutant cerebellum (Fig. 2 and Fig. S2). Indeed, oxidative phosphorylation and pathways/GO related to mitochondria and ribosomes, all known to support the generation of ROS, were enriched in mutant animals. Furthermore, cell deconvolution analysis identified the microglia as an important player in the changes described in the *Cntnap2^−/−^* cerebellum (Fig. 2 and Fig. S3). Importantly, C4b, C1qtnf1, and C3 complement genes were also less expressed in the mutant cerebellum. As the complement system has been shown to mediate the interaction between microglia cells and neurons (32), impaired communication between these two cell types may be present in the *Cntnap2^−/−^* cerebellum as a consequence of dysfunctional complement system. In parallel, pro-inflammatory changes in the PB of *Cntnap2^−/−^*mice were described. CNTNAP2 is involved in the clustering and localization of voltage-gated potassium (Kv) channels (33). Whether these pro-inflammatory changes are linked to altered cerebellar excitability (30,31) and localization and functions of Kv channels remains to be investigated.

To target oxidative stress in the cerebellum, we administered 50 mg/kg NAC i.p. for 28 days to *Cntnap2^−/−^* and Cntnap2^+/+^ adult mice. NAC is an antioxidant molecule known to scavenge ROS in glutathione-dependent and independent manner (34). While hyperactivity and social impairments were completely rescued in mutant animals, we unexpectedly observed that social deficits were induced in *Cntnap2^+/+^* animals treated with NAC (Fig. 4). These results were mirrored by the molecular changes observed in the cerebellum. Indeed, while levels of ROS and pro-inflammatory molecules in mutant mice were decreased by NAC treatment, neuronal oxidative stress and levels of pro-inflammatory molecules in the cerebellum were increased in NAC-treated *Cntnap2^+/+^*mice (Fig. 5). It is commonly accepted that oxidative stress is induced when ROS levels exceed the amount of ROS-scavenging (antioxidant) molecules. Despite this, the equilibrium may be broken even when levels of antioxidant molecules are higher than ROS, and thus the so-called “antioxidative stress” is induced (35,36). As *Cntnap2^+/+^* mice display low ROS levels, we can therefore assume that the administration of NAC may induce antioxidative stress in the cerebellum of these mice. In addition, as low levels of ROS are required for physiological processes (37), a condition of stress is induced when oxygen radicals are completely removed. For this reason, NAC concentrations must be cautiously titrated to avoid side effects.

Previous studies showed that ASD-related behaviors are induced after the administration of microglia-secreted TNF and reverted by microglia depletion and TNF inhibition (38). In line with this study, we described that a link may exist between microglia dysfunction and ASD-related behaviors. Indeed, microglia branching in Crus1/2 cerebellar areas was consistently improved in NAC-treated mutant mice (Fig. 6). The observation that NAC can suppress microglial inflammation via TNF signaling (39) indicates a possible mechanism of action occurring in the cerebellum of NAC-treated *Cntnap2^−/−^* mice. Conversely, both frequency and branching of microglia cells were impaired in *Cntnap2^+/+^* animals after NAC administration (Fig. 6). Of note, Crus1/2 cerebellar regions are known to play a key role in social behaviors (21–23). While reduced levels of IL-1β, TNF, IL-6, CCL3, MMP12, and IL-22 and increased levels of SOD3 were found within cerebellar microglia cells from NAC-treated mutant animals, increased levels of almost all molecules and decreased levels of SOD3 were present in the cerebellum of NAC-treated *Cntnap2^+/+^* mice.

Similar results were found in the periphery. Indeed, decreased ROS levels and a lower expression of IL-6, TNF and IFNγ were detected in the PB, bone marrow, and spleen of NAC-treated *Cntnap2^−/−^* mice (Fig. 7). Although NAC treatment induced the synthesis of glutathione in *Cntnap2^+/+^*mice (as indicated by the increased levels of cys-gly in the serum), the overall ROS levels were higher when compared to control mice (Fig. 7). Despite this, the expression of pro-inflammatory molecules in NAC-treated *Cntnap2^+/+^* mice did not change as observed in the cerebellum. Although the reason of this finding is still unknown, we can speculate that in NAC-treated *Cntnap2^+/+^* mice the mechanisms to control the onset of pro-inflammatory processes are still effective in the periphery, while they may be impaired in the cerebellum.

Importantly, many of the pro-inflammatory changes detected in *Cntnap2^−/−^*mice were found within CD14^+^ cells. While CD14 in the PB is a monocyte marker and in the bone marrow and spleen it defines monocytes and macrophages, this molecule is expressed by activated microglia and macrophages in the cerebellum (40,41). As Tmem119 is lost after microglia cell activation (42) (in our settings after PMA/Ionomycin stimulation; see Materials and Methods), we used CD14 as a microglia marker in flow cytometry. Although still controversial, it has been suggested that blood monocytes may differentiate into microglia cells, particularly in the presence of pro-inflammatory conditions (43). Indeed, monocytes are known to cross the blood-brain barrier, enter the brain and differentiate into microglia-like cell types. We can therefore hypothesize that a strong connection may exist between monocytes present in the blood and microglia cells within the cerebellum.

Taken together, our findings suggest that the interplay between oxidative stress and pro-inflammatory conditions in the cerebellum may support ASD-related behaviors. A connection was found between cerebellar inflammation and levels of pro-inflammatory molecules in the PB. Further studies are needed to understand whether cerebellar inflammation may be induced by immune cells migrating into the brain from the periphery, or alternatively whether pro-inflammatory conditions may be induced within the cerebellum and afterwards be “exported” to the periphery by circulating immune cells.

## Materials and Methods

### Animals

All experimental procedures were approved by the Animal Welfare Committee of the University of Trento and Italian Ministry of Health (protocol n.922/2018-PR and n. 547/2021-PR) according to the European Community Directive 2010/63/EU. Mice were housed in a 12 h light/dark cycle with food and water available *ad libitum*, taking care to minimize animal’s pain and discomfort. Male and female *Cntnap2^+/+^* and *Cntnap2^−/−^* age-matched adult littermates (6–9 months old; weight 25–35 g) obtained from heterozygous matings were used. Numbers of mice used for each experiment are shown in the Figure legends.

### Tissue harvesting

Brain tissues used for RT-PCR were dissected and frozen in dry ice. Brain samples used for flow cytometry were homogenized with pestles immediately after dissections and single-cell suspension were generated using Falcon 70 μm cell strainer (Corning). Two hundred μl of PB was harvested from each mouse and collected in a heparinized tube. PBMCs were isolated using a Ficoll-Hypaque density gradient (Sigma-Aldrich). BM cells were isolated by flushing femurs and tibias with PBS. Spleen samples were smashed through Falcon 70 µm cell strainers. After the isolation, PBMCs, BM and spleen cells were washed once with RPMI 1640 (Sigma-Aldrich) and resuspended in complete medium (RPMI 1640 supplemented with 10% fetal calf serum, FCS, 100 U/mL penicillin and 100 μg/mL streptomycin; Sigma-Aldrich and Invitrogen respectively). After the isolation, brain cells were washed with complete medium and incubated for 12h at 37°C.

### RNA isolation and quantitative RT-PCR (qRT-PCR)

Total RNAs were extracted from cerebral cortex, hippocampus and cerebella of adult Cntnap2^+/+^ and Cntnap2^−/−^ mice mice using RNeasi Plus Mini Kit (Qiagen), and retro-transcribed to cDNA according to published protocols (44). qRT-PCR was performed in a Bio-Rad C1000 Thermal Cycler, using the PowerUp™ SYBR™ Green Master Mix (Applied Biosystems). Primers (Eurofins Genomics) were designed on different exons to avoid amplification of genomic DNA. Sequences of primers using for the study are shown in TableS6. CFX3 Manager 3.0 (Bio-Rad) software was used to perform expression analyses. Mean cycle threshold (Ct) values from triplicate experiments were calculated for each gene of interest and the housekeeping gene β actin, and then corrected for PCR efficiency and inter-run calibration. The expression level of each mRNA of interest (normalized against β actin) was compared from triplicate experiments performed on RNA pools from 8-10 samples per group.

### RNAseq analysis

#### Preparation of samples

RNA sequencing was performed on the RNAs extracted from cerebella of 8 adult *Cntnap2^+/+^* (4 females, 4 males) and 8 *Cntnap2^−/−^* (4 females, 4 males). To determine the quality and the integrity of the RNA samples, RNA Integrity Numbers (RIN) were assessed using a 2100 Bioanalyzer Instrument (Agilent). Only samples with RIN≥ 8 were considered for the analysis. Libraries were prepared using a TruSeq® Stranded mRNA Library Prep (48 Samples) Kit (Illumina). Library preparation, quality check, and sequencing were carried out using Illumina NovaSeq6000 technology by Next Generation Sequencing Core Facility, at the Department of Cellular, Computational and Integrative Biomedicine (CIBIO) of the University of Trento (Italy).

#### Differential gene expression and gene set enrichment analysis

After generating the gene count matrix, we discarded the genes with count-per-million lower than 100 in 20% of the samples and used the resulting filtered count matrix as an input for calling differentially expressed genes (DEGs) using the R package DESeq2 (1.34.0) (45). The counts were normalized for differences in sequencing depth of the samples (“median ratio method” described by Equation 5 in Anders and Huber; 46) and for each gene, dispersion values were calculated under the assumption of a negative binomial distribution (47). Negative binomial generalized linear models were fitted to the normalized count matrix including the genotype of the mice (our variable of interest) as covariate. Wald test was then performed for each gene to determine whether it was differentially expressed between the two genotypes and the resulting P-value was corrected using the Benjamini-Hochberg (BH) adjustment. lfcShrink function of the apeglm R package (1.16.0) was additionally performed to shrink the log2 fold changes as these estimates may not represent true differences in expression due to high variance in read counts. For downstream analyses, we used the regularized-logarithm transformation (46) on the count data to stabilize the variance across the mean. Gene set enrichment analysis (GSEA) was performed on over- and under-represented genes included in the DEGs. Both gene ontology (GO) and KEGG pathway analyses were performed on the genes.

#### Cell deconvolution analysis

Relative proportions of cell-types were inferred with the algorithm MuSiC using the single-nuclei RNAseq signature matrix generated by Kozareva *et al.* (19). The signature matrix included 24,409 genes and 200 replicates for 18 cerebellar cell types. Non-parametric multivariate analysis of variance (NPMANOVA) was performed to test if there was a significant difference in the global fraction of cells between genotypes. Bonferroni post-hoc test was used to determine which group contributed the most to the variance in global cell fraction. Finally, Kruskal-Wallis test was performed to assess differences between genotypes for each cell-type proportion.

### NAC treatment and behavioural tests

*Cntnap2^+/+^* and *Cntnap2^−/−^* mice were injected intraperitoneally with 50mg/kg NAC (Sigma-Aldrich) dissolved in PBS or vehicle (PBS) for 28 consecutive days. Optimal NAC concentration to reduce cerebellar inflammation in Cntnap2^−/−^ mice was assessed based on the available literature (48) and tested in preliminary experiments performed on *Cntnap2^−/−^* mice (data not shown).

#### Behavioral tests

Open field, rotarod and 3-chamber social tests were performed during the last 5 days of NAC treatment of *Cntnap2^+/+^* and *Cntnap2^−/−^* mice. For all behavioral tests, male and female mice were habituated and tested separately to avoid experimental noise.

#### Open field test

Open field test was performed to assess motor activity of mice. Animals were placed in empty open field arena (40 cm × 40 cm × 40 cm) and allowed to freely explore it for 10 min. Sessions were recorded by an overhead camera placed over the arena and mice were automatically video tracked using the software EthoVisionXT (Noldus). Mean velocity and distance moved in the arena were analyzed.

#### Rotarod test

To measure cerebellar-associated motor coordination, the rotarod test was performed. A habituation phase conducted at constant speed of 4 rpm was performed in the two days before the test. Experimental phase consisted in 3 trials in which the rotation speed was increased from 4 rpm to 64 rpm. Falling mice landed on a metallic platform that was connected to a timer reporting the time spent on the rotating rod. The average time spent on the rod (latency to fall) and the average RPM of the three trials were calculated and used as quantitative indicators of the motor ability of the mice to stay balanced on the rotating accelerating rod.

#### Three-Chamber social test

The three-Chamber social test was used to assess the social behaviors of mice (49). The apparatus consisted of a plexiglass rectangular box (60×40×22(h) cm, each chamber 20×40×22(h) cm) having grayed colored walls and with removable panels separating the box into three chambers. On the four days before the beginning of the experimental phase, mice underwent a habituation phase in which they were placed in the three-chamber apparatus and allowed to freely explore it for 10 min. The experimental phase consisted of a 5-minutes habituation session, a 10-minutes exploration session and the sociability test, in which one unfamiliar mouse was placed into a wire cylindrical cage (20 cm in height, 10 cm bottom-diameter, 1cm bars spaced) in one of the up corner of an external chamber. An identical empty wire cage was placed in the opposite external chamber. The tested mouse was allowed to freely interact with the unfamiliar mouse and with wire cages for 10 minutes. The time spent in the chambers, as well as the sociability index (the ration between the time spent in the social chamber and the total time spent in the external chambers) was measured to assess sociability of the tested mouse. Trials were recorded by an overhead camera placed over the three-chamber apparatus. Mice were automatically video tracked using EthoVisionXT.

### Flow cytometry and FACS sorting

Flow cytometry was used to measure cytokine levels and immune cell populations in the cerebellum, PB, BM and spleen. Immunofluorescence surface staining was performed by adding a panel of directly conjugated antibody to freshly prepared cells. Dead cells were excluded from the analysis using fixable viability dye (FVD) BV421 (BD Biosciences). To assess the expression of cytokines, cells were incubated with 30 ng/mL phorbol 12-myristate 13-acetate (PMA) and 500 ng/mL ionomycin in the presence of 10 mg/mL brefeldin A (BFA) (all molecules from Sigma-Aldrich) for 4 h at 37 °C. After surface staining, cells were permeabilized using the Cytofix/Cytoperm kit (BD Biosciences), and incubated with intracellular antibodies. Labeled cells were measured using a FACS Symphony 1 (BD Biosciences). Data were analyzed using Flowjo software. The antibodies used in the experiments are shown in TableS7. ROS levels were measured after incubation of cerebellum cells and PBMCs with the fluorescent dye dihydroethidium (Sigma-Aldrich) at a concentration of 1:250 in complete medium for 20 min at 37 °C.

Fluorescence-activated cell sorting (FACS) of microglia cells and neurons was performed using cerebellar samples from NAC-treated and untreated Cntnap2^+/+^ and Cntnap2^−/−^ mice. For each experimental group 5 samples were pooled. Single-cell suspensions were incubated for 20 minutes with Tmem119 and Neun antibodies and afterwards sorted with the FACS sorter Aria Fusion (BD Biosciences) at the Institute for Biomedical Aging Research, University of Innsbruck.

### Thiols determination

Serum Cysteine (Cys), Cysteinyl-Glycine (Cys-Gly), and Glutathione (GSH) levels were analyzed as previously reported (50)

### Immunofluorescence staining of cerebellar sections

After NAC treatment, brains were excised, post-fixed overnight in 4% paraformaldehyde (PFA) at 4°C, washed ans switched to a cryoprotectant solution (80% PBS, 20% glycerol with 0.1% sodium azide) and stored at 4°C. Cryoprotectant brains were then included in 4% agarose gel and sectioned sagitally on a vibratome (Leica, VT1200) at 40μm thickness. Sequential slices were collected in separate wells containing cryoprotectant solution.

Free-floating slices were firstly rinsed in PBS and then washed three times in PBS containing 0.2% Triton-X (Fisher, AC215680010) for 10 minutes each. Tissue slices were then incubated in blocking solution (10% FCS, 0.2% Triton in PBS) for 1 hour and subsequentially co-incubated overnight at room temperature in primary antibody solution (rabbit Iba-1, 1:1000, and mouse Tmem119, 1:500, both Synaptic Systems) prepared in blocking solution. This step was followed by 2 hours incubation in Alexa Fluor anti-rabbit 488 (1:350, Life Technologies) and horse anti-mouse biotinylated (1:350 Life Technologies). Sections were then incubated in streptavidin-AF594 solution (1:750 μl, Life Technologies) for 1 hour at room temperature. Tissue sections were mounted on slides, dried for 10 minutes and cover-slipped with fluorescent mounting medium (Southern biotech). Slides were then stored at 4°C until use.

### Image acquisition with confocal microscope

A confocal laser scanning microscope Leica TCS-SP8, equipped with a HC PL CS2 40X objective and interfaced with Leica “LAS-X” software was used to scan the entire area through the full *x*-, *y*-, and *z*-axes of Crus1 and Crus2 in the cerebellum. Images were acquired at the resolution of 1024×1024 and 400 Hz scan speed. Excitation/emission wavelengths were 493/537 for Alexa-488 and 600/670 for Alexa-594 fluorophore. Acquisition parameters were set during the first acquisition of control experimental sample (*Cntnap2^+/+^* PBS) and kept consistent for all the images acquisition. All acquisitions were anonymized to avoid bias during acquisition and data analysis.

#### Analysis of confocal microscope images

Images were analyzed using Fiji-ImageJ software. Images were pre-processed to allow a more consistent and reliable analysis throughout the experimental conditions. Firstly, an interval of 80 stacks (corresponding to the total 40µm thickness of each sections) was set for each image, then the background was subtracted to allow further steps of analysis. Quantification of microglia cell numbers and Sholl analysis within Crus1 and Crus2 areas were performed on Iba-1^+^ cells.

#### Quantification of microglia cells

Borders of molecular and granular layers were traced, and the areas were measured using the ImageJ ROI manager tool. Soma of Iba-1^+^ cells were manually counted using cell-counter plugin. The total quantification of iba-1^+^ cells in each layer and in the total area (the sum of molecular and granular layers) was given by the ratio of the number of somas/area(0.1mm^2^).

#### Sholl analysis of microglia cells

Iba-1^+^ cells were randomly selected in the molecular and granular layer, respectively 1 cell in each layer for each section. Cells were extracted, then segmented and tracked using the ImageJ tool “microglia plugin”. The resulting output was threshold and extra pixels adjusted with “erode/dilate tool”. Sholl analysis was then performed using the “Sholl Analysis plugin”. The number of total intersections were calculated as the sum of intersections at each concentric shells of analysis.

### Statistics

Statistical analyses were performed with GraphPad Prism 8.0 software, using two-way ANOVA followed by Tukey post-hoc test, unpaired t-tests, and Kruskal-Wallis test with the level of significance set at p<0.05.

---

**Figure S1.**
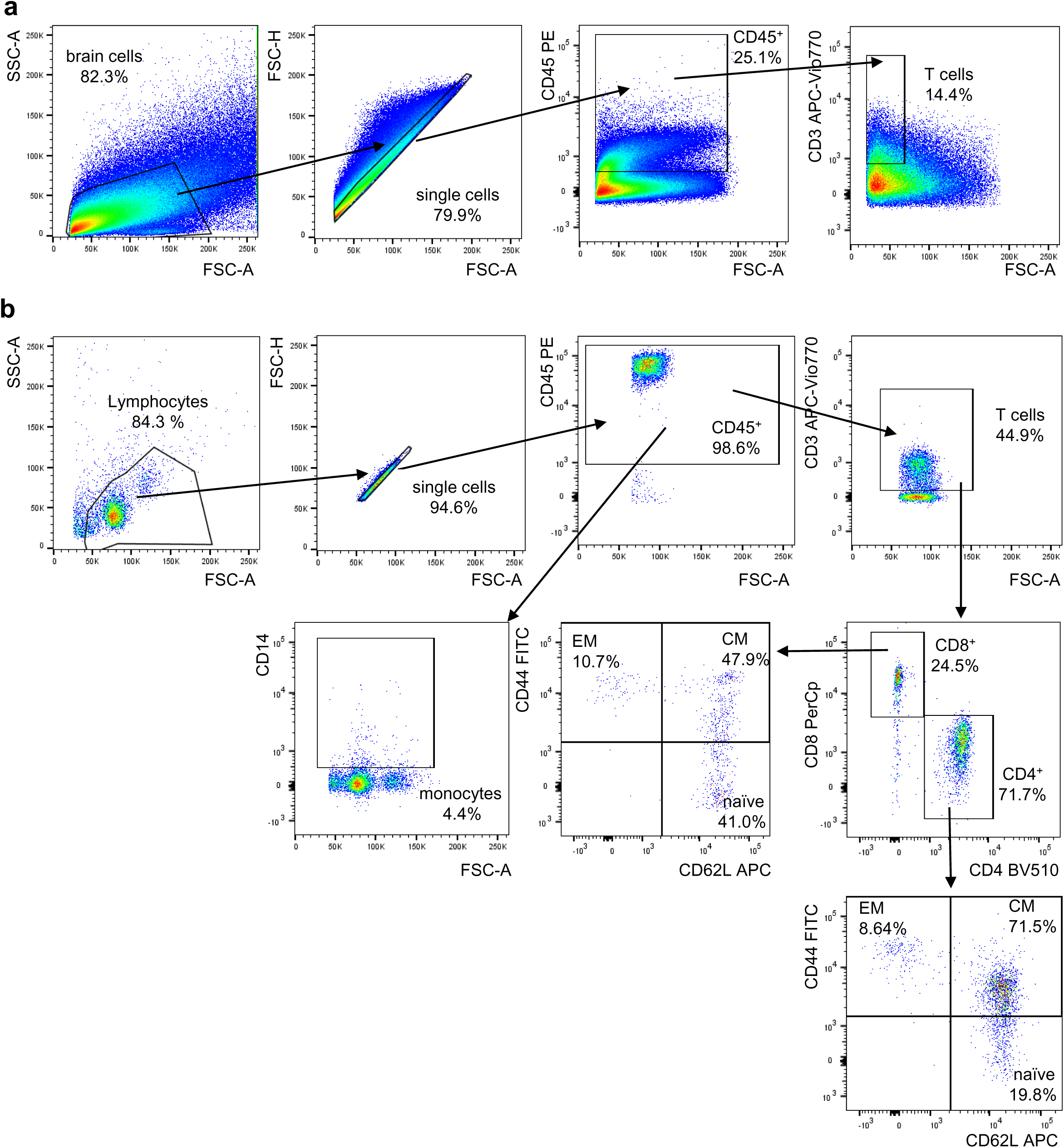
Gating strategy used for the flow cytometry experiments. **(a)** Gating strategy used to define (**a**) CD45^+^ cells and T cells within brain cells, and (**b**) CD45^+^ cells, monocytes (CD14^+^ cells), T cells (CD8^+^ and CD4^+^) and CD44^+^CD62L^−^ EM, CD44^+^CD62L^+^ CM and CD44^−^CD62L^+^ naïve cells in CD8^+^ and CD4^+^ T cells within PBMCs.

**Figure S2.**
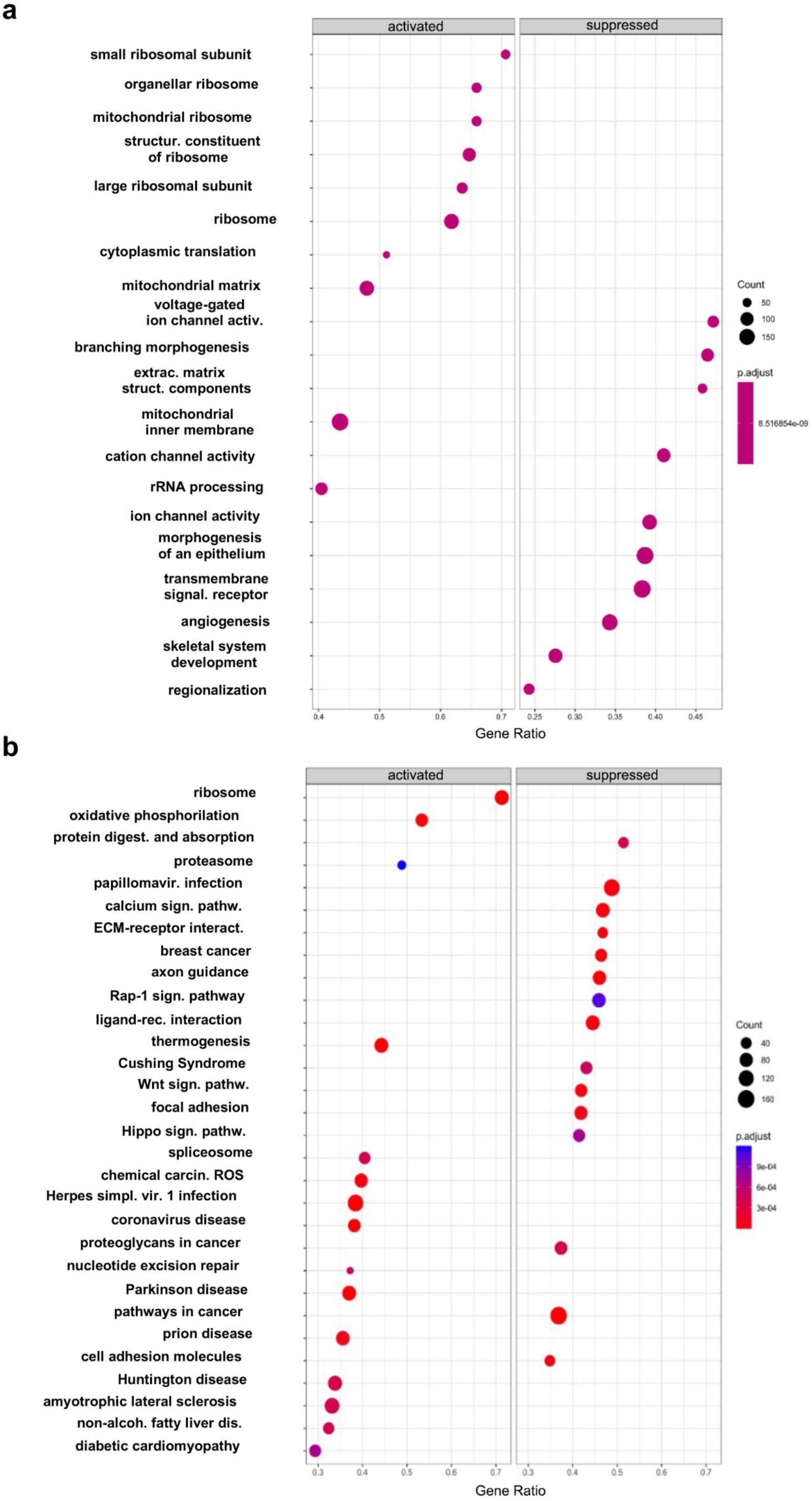
GO and KEGG pathway analysis of RNAseq data. Positively and negatively enriched (a) GO and (b) KEGG pathways obtained after the GSEA.

**Figure S3.**
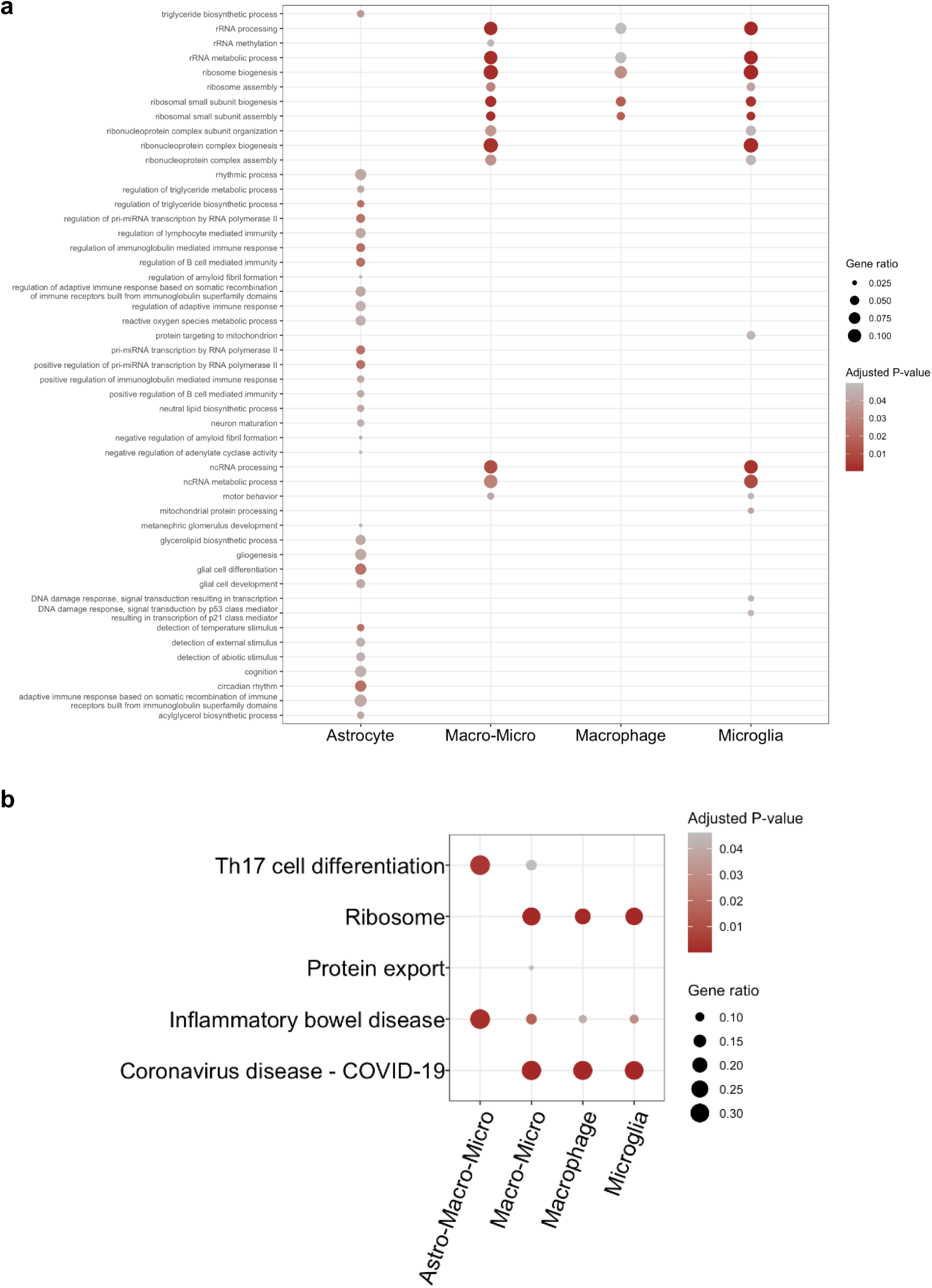
GO and KEGG pathway analysis of RNAseq data after cell deconvolution analysis. (**a**) GO and (**b**) KEGG pathway analysis after adjusting for cell subsets (astrocytes, macrophages+microglia, macrophages, and microglia. Adjusted p values are displayed in the graphs using the scale color.

**Figure S4.**
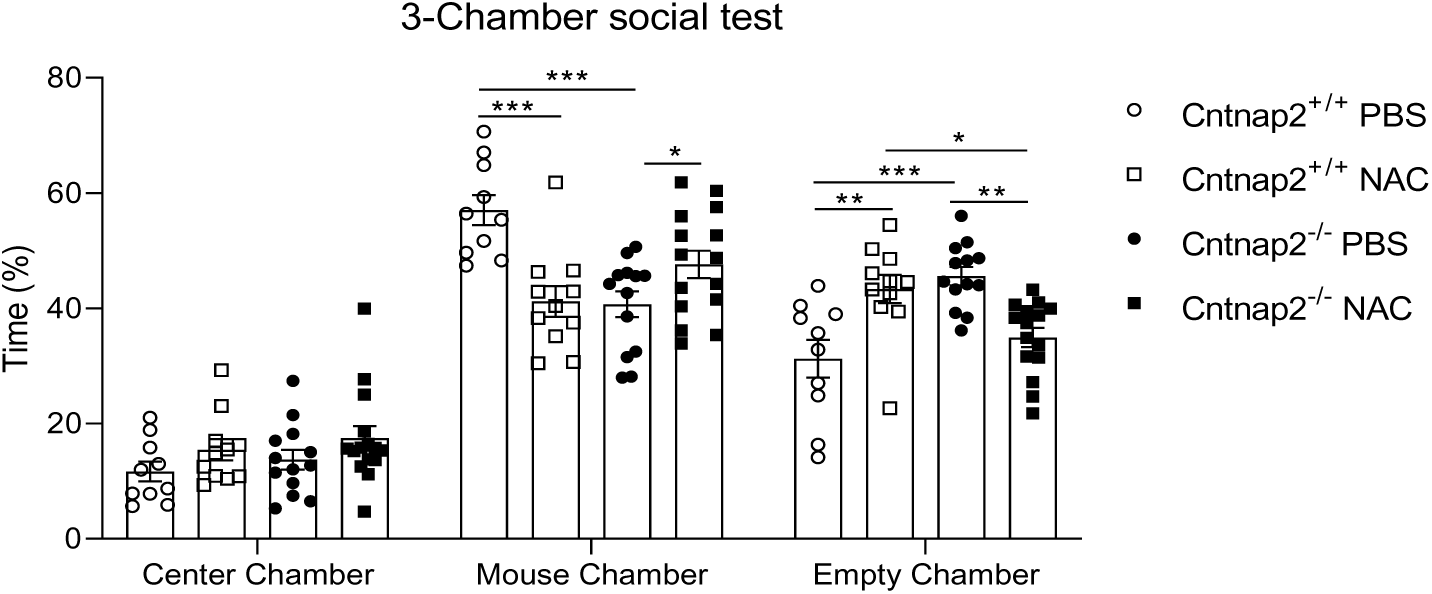
Percentage of time in the 3 chambers in the 3-chamber test. Percentage of time in the center chamber, mouse chamber and empty chamber in the 3-chamber social test. Two-way ANOVA, Tukey post-hoc test. *p<0.05; **p<0.01; ***p<0.001. n_Cntnap2+/+PBS=_7, n_Cntnap2+/+NAC=_6, n_Cntnap2−/−PBS=_7, n_Cntnap2+/+NAC=_8. Two-way ANOVA, Tukey post-hoc test. *p<0.05; ****p<0.0001.

**Figure S5.**
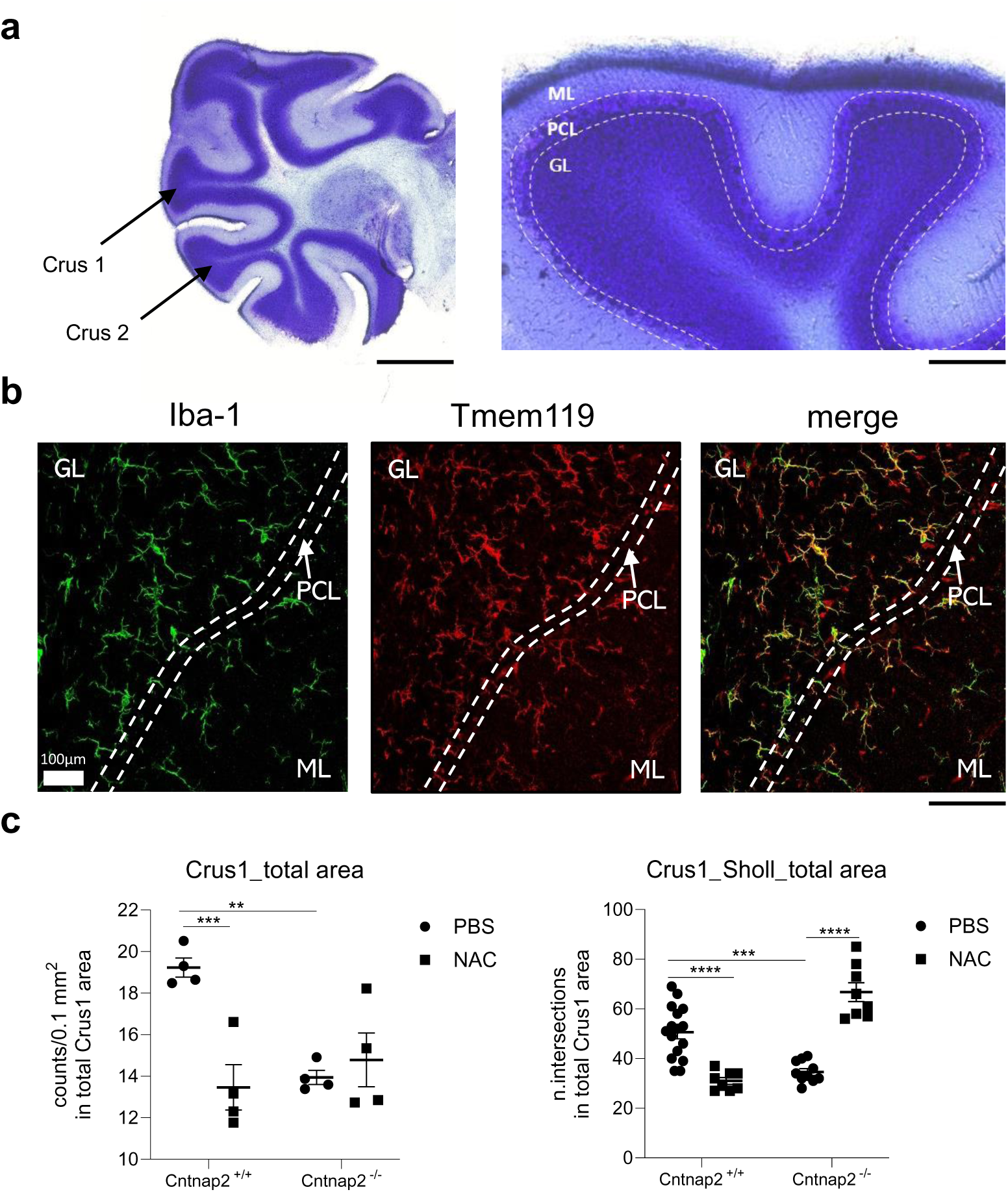
Analysis of microglia cells in the cerebellum of PBS- and NAC-treated mice. (**a**) representative picture of Nissl staining in the cerebellum of *Cntnap2^+/+^* mice. Crus1 and Crus 2 areas, with their respective molecular layer (ML), Purkinje cells layer (PCL) and granular layer (GL) are displayed. (**b**) Representative images showing the colocalization between iba-1^+^ and tmem119^+^ cells within Crus2 area. (**c**) iba-1^+^ cells (expressed in 0.1 mm2) and (**d**) intersections calculated with the Sholl analysis in the Crus1 total area. For cell number quantification: n=4 in each group; For Sholl analysis: n_Cntnap2+/+PBS=_16, n_Cntnap2+/+NAC=_8, n_Cntnap2−/−PBS=_10, n_Cntnap2+/+NAC=_8. Two-way ANOVA, Tukey post-hoc test. *p<0.05; **p<0.01; ***p<0.001; ****p<0.0001.

**Figure S6.**
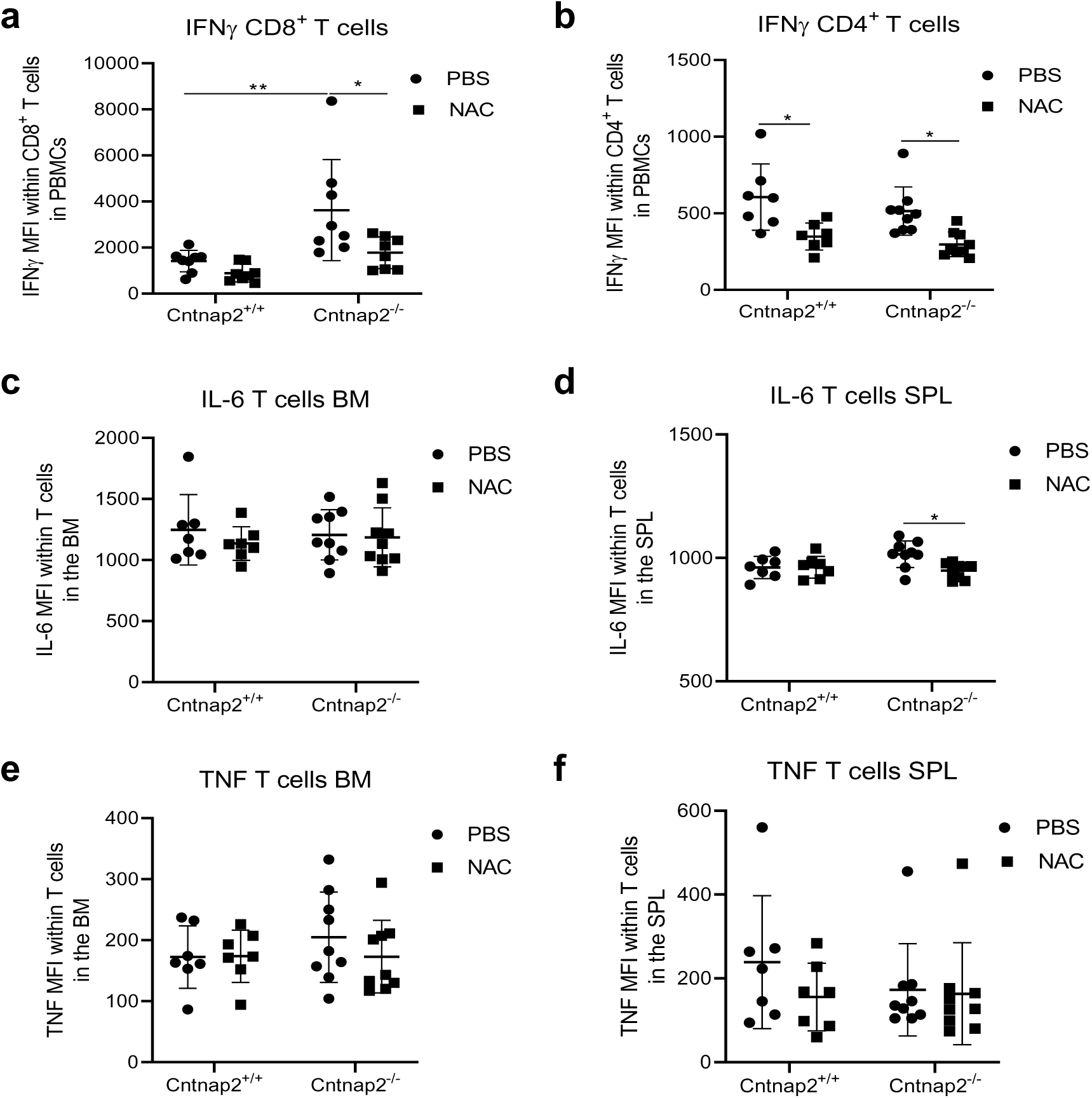
The impact of NAC on pro-inflammatory molecules within T cell subsets in the peripheral blood, bone marrow, and spleen of PBS- and NAC-treated *Cntnap2^+/+^* and *Cntnap2^−/−^* mice. Mean fluorescence intensity (MFI) of (a) IFNγ in CD8^+^ T cells, and (b) IFNγ in CD4^+^ T cells within PBMCs. MFI of IL-6 in CD8^+^ T cells within (c) the bone marrow, and (d) the spleen. MFI of TNF in CD8^+^ T cells within (e) the bone marrow, and (f) the spleen. n=7-9 in each group. Two-way ANOVA, Tukey post-hoc test. *p<0.05; **p<0.01.

## References

1. American Psychiatric Association. Diagnostic and Statistical Manual of Mental Disorders, 5th ed.; American Psychiatric Publishing: Washington, DC, USA, 2013.

2. Zeidan J, Fombonne E, Scorah J, Ibrahim A, Durkin MS, Saxena S, Yusuf A, Shih A, Elsabbagh M. Global prevalence of autism: A systematic review update. Autism Res. 2022; 15(5):778–790. doi: 10.1002/aur.2696.

3. Pangrazzi L, Balasco L, Bozzi Y. Oxidative Stress and Immune System Dysfunction in Autism Spectrum Disorders. Int J Mol Sci. 2020 May 6;21(9):3293. doi: 10.3390/ijms21093293

4. Ashwood, P.; Krakowiak, P.; Hertz-Picciotto, I.; Hansen, R.; Pessah, I.; Van De Water, J. Elevated plasma cytokines in autism spectrum disorders provide evidence of immune dysfunction and are associated with impaired behavioral outcome. Brain, Behav. Immun. 2010, 25, 40–45. doi: 10.1016/j.bbi.2010.08.003

5. Inga Jácome MC, Morales Chacòn LM, Vera Cuesta H, Maragoto Rizo C, Whilby Santiesteban M, Ramos Hernandez L, Noris García E, González Fraguela ME, Fernandez Verdecia CI, Vegas Hurtado Y, Siniscalco D, Gonçalves CA, Robinson-Agramonte ML. Peripheral Inflammatory Markers Contributing to Comorbidities in Autism. Behav Sci (Basel). 2016; 6(4):29. doi: 10.3390/bs6040029

6. Li X, Chauhan A, Sheikh AM, Patil S, Chauhan V, Li XM, Ji L, Brown T, Malik M. Elevated immune response in the brain of autistic patients. J Neuroimmunol. 2009; 207(1-2):111–6. doi: 10.1016/j.jneuroim.2008.12.002

7. Vargas DL, Nascimbene C, Krishnan C, Zimmerman AW, Pardo CA. Neuroglial activation and neuroinflammation in the brain of patients with autism. Ann Neurol. 2005; 57(1):67–81. doi: 10.1002/ana.20315. Erratum in: Ann Neurol. 2005 Feb;57(2):304

8. Chen L, Shi XJ, Liu H, Mao X, Gui LN, Wang H, Cheng Y. Oxidative stress marker aberrations in children with autism spectrum disorder: a systematic review and meta-analysis of 87 studies (N = 9109). Transl Psychiatry. 2021; 11(1):15. doi: 10.1038/s41398-020-01135-3

9. James SJ, Melnyk S, Jernigan S, Cleves MA, Halsted CH, Wong DH, Cutler P, Bock K, Boris M, Bradstreet JJ, Baker SM, Gaylor DW. Metabolic endophenotype and related genotypes are associated with oxidative stress in children with autism. Am J Med Genet B Neuropsychiatr Genet. 2006; 141B(8):947-56. doi: 10.1002/ajmg.b.30366

10. Rose S, Melnyk S, Pavliv O, Bai S, Nick TG, Frye RE, James SJ. Evidence of oxidative damage and inflammation associated with low glutathione redox status in the autism brain. Transl Psychiatry. 2012;2(7):e134. doi: 10.1038/tp.2012.61

11. Liu X, Lin J, Zhang H, Khan NU, Zhang J, Tang X, Cao X, Shen L. Oxidative Stress in Autism Spectrum Disorder-Current Progress of Mechanisms and Biomarkers. Front Psychiatry. 2022; 13:813304. doi: 10.3389/fpsyt.2022.813304

12. Ghezzo A, Visconti P, Abruzzo PM, Bolotta A, Ferreri C, Gobbi G, Malisardi G, Manfredini S, Marini M, Nanetti L, Pipitone E, Raffaelli F, Resca F, Vignini A, Mazzanti L. Oxidative Stress and Erythrocyte Membrane Alterations in Children with Autism: Correlation with Clinical Features. PLoS One. 2013; 8(6):e66418. doi: 10.1371/journal.pone.0066418

13. Popa-Wagner A, Mitran S, Sivanesan S, Chang E, Buga AM. ROS and brain diseases: the good, the bad, and the ugly. Oxid Med Cell Longev. 2013;2013:963520. doi: 10.1155/2013/963520.

14. Pangrazzi L, Meryk A, Naismith E, Koziel R, Lair J, Krismer M, Trieb K, Grubeck-Loebenstein B. “Inflamm-aging” influences immune cell survival factors in human bone marrow. Eur J Immunol. 2017; 47(3):481–492. doi: 10.1002/eji.201646570

15. Peñagarikano O, Abrahams BS, Herman EI, Winden KD, Gdalyahu A, Dong H, Sonnenblick LI, Gruver R, Almajano J, Bragin A, Golshani P, Trachtenberg JT, Peles E, Geschwind DH. Absence of CNTNAP2 leads to epilepsy, neuronal migration abnormalities, and core autism-related deficits. Cell. 2011;147(1):235–46. doi: 10.1016/j.cell.2011.08.040.

16. Vogt D, Cho KKA, Shelton SM, Paul A, Huang ZJ, Sohal VS, Rubenstein JLR. Mouse Cntnap2 and Human CNTNAP2 ASD Alleles Cell Autonomously Regulate PV+ Cortical Interneurons. Cereb Cortex. 2018; 28(11):3868–3879. doi: 10.1093/cercor/bhx248

17. Smolders J, Heutinck KM, Fransen NL, Remmerswaal EBM, Hombrink P, Ten Berge IJM, van Lier RAW, Huitinga I, Hamann J. Tissue-resident memory T cells populate the human brain. Nat Commun. 2018;9(1):4593. doi: 10.1038/s41467-018-07053-9

18. Hughes CE, Nibbs RJB. A guide to chemokines and their receptors. FEBS J. 2018;285(16):2944–2971. doi: 10.1111/febs.14466

19. Kozareva V, Martin C, Osorno T, Rudolph S, Guo C, Vanderburg C, Nadaf N, Regev A, Regehr WG, Macosko E. A transcriptomic atlas of mouse cerebellar cortex comprehensively defines cell types. Nature. 2021;598(7879):214–219. doi: 10.1038/s41586-021-03220-z

20. Tardiolo G, Bramanti P, Mazzon E. Overview on the Effects of N-Acetylcysteine in Neurodegenerative Diseases. Molecules. 2018 13;23(12):3305. doi: 10.3390/molecules23123305

21. Heleven E, van Dun K, Van Overwalle F. The posterior Cerebellum is involved in constructing Social Action Sequences: An fMRI Study. Sci Rep. 2019; 9(1):11110. doi: 10.1038/s41598-019-46962-7

22. Stoodley CJ, D’Mello AM, Ellegood J, Jakkamsetti V, Liu P, Nebel MB, GibsonJM, Kelly E, Meng F, Cano CA, Pascual JM, Mostofsky SH, Lerch JP, Tsai PT. Altered cerebellar connectivity in autism and cerebellar-mediated rescue of autism-related behaviors in mice. Nat Neurosci. 2017;20(12):1744–1751. doi: 10.1038/s41593-017-0004-1

23. Kelly E, Meng F, Fujita H, Morgado F, Kazemi Y, Rice LC, Ren C, Escamilla CO, Gibson JM, Sajadi S, Pendry RJ, Tan T, Ellegood J, Basson MA, Blakely RD, Dindot SV, Golzio C, Hahn MK, Katsanis N, Robins DM, Silverman JL, Singh KK, Wevrick R, Taylor MJ, Hammill C, Anagnostou E, Pfeiffer BE, Stoodley CJ, Lerch JP, du Lac S, Tsai PT. Regulation of autism-relevant behaviors by cerebellar-prefrontal cortical circuits. Nat Neurosci. 2020;23(9):1102–1110. doi:10.1038/s41593-020-0665-z.

24. Rylaarsdam L, Guemez-Gamboa A. Genetic Causes and Modifiers of Autism Spectrum Disorder. Front Cell Neurosci. 2019 Aug 20;13:385. doi: 10.3389/fncel.2019.00385

25. Sydnor LM, Aldinger KA. Structure, Function, and Genetics of the Cerebellum in Autism. J Psychiatr Brain Sci. 2022;7:e220008. doi: 10.20900/jpbs.20220008

26. Tan GC, Doke TF, Ashburner J, Wood NW, Frackowiak RS (2010) Normal variation in fronto-occipital circuitry and cerebellar structure with an autism-associated polymorphism of CNTNAP2. Neuroimage 53:1030–1042. doi: 10.1016/j.neuroimage.2010.02.018.

27. Becker EB, Zuliani L, Pettingill R, Lang B, Waters P, Dulneva A, Sobott F, Wardle M, Graus F, Bataller L, Robertson NP, Vincent A (2012) Contactin-associated protein-2 antibodies in non-paraneoplastic cerebellar ataxia. J Neurol Neurosurg Psychiatry 83:437–440. doi: 10.1136/jnnp-2011-301506.

28. Melzer N, Golombeck KS, Gross CC, Meuth SG, Wiendl H (2012) Cytotoxic CD81 T cells and CD1381 plasma cells prevail in cerebrospinal fluid in non-paraneoplastic cerebellar ataxia with contactin-associated protein-2 antibodies. J Neuroinflammation 9:160. doi: 10.1186/1742-2094-9-160.

29. Tsai PT. Autism and cerebellar dysfunction: Evidence from animal models. Semin Fetal Neonatal Med. 2016;21(5):349–55. doi: 10.1016/j.siny.2016.04.009

30. Kloth AD, Badura A, Li A, Cherskov A, Connolly SG, Giovannucci A, Bangash MA, Grasselli G, Peñagarikano O, Piochon C, Tsai PT, Geschwind DH, Hansel C, Sahin M, Takumi T, Worley PF, Wang SS. Cerebellar associative sensory learning defects in five mouse autism models. Elife. 2015;4:e06085. doi: 10.7554/eLife.06085.

31. Fernández M, Sánchez-León CA, Llorente J, Sierra-Arregui T, Knafo S, Márquez-Ruiz J, Peñagarikano O. Altered Cerebellar Response to Somatosensory Stimuli in the Cntnap2 Mouse Model of Autism. eNeuro. 2021 Oct 21;8(5):ENEURO.0333-21.2021. doi: 10.1523/ENEURO.0333-21.2021.

32. Schafer DP, Lehrman EK, Kautzman AG, Koyama R, Mardinly AR, Yamasaki R, Ransohoff RM, Greenberg ME, Barres BA, Stevens B. Microglia sculpt postnatal neural circuits in an activity and complement-dependent manner. Neuron. 2012;74(4):691–705. doi: 10.1016/j.neuron.2012.03.026

33. Saint-Martin M, Joubert B, Pellier-Monnin V, Pascual O, Noraz N, Honnorat J. Contactin-associated protein-like 2, a protein of the neurexin family involved in several human diseases. Eur J Neurosci. 2018;48(3):1906–1923. doi: 10.1111/ejn.14081.

34. Kannan N, Nguyen LV, Makarem M, Dong Y, Shih K, Eirew P, Raouf A, Emerman JT, Eaves CJ. Glutathione-dependent and -independent oxidative stress-control mechanisms distinguish normal human mammary epithelial cell subsets. Proc Natl Acad Sci U S A. 2014, 111(21):7789–94. doi: 10.1073/pnas.1403813111

35. Villanueva C, Kross RD. Antioxidant-induced stress. Int J Mol Sci. 2012;13(2):2091–2109. doi: 10.3390/ijms13022091

36. Dündar Y, Aslan R. Antioxidative stress. Eastern J. Med. 2000;5:45–47

37. Sena LA, Chandel NS. Physiological roles of mitochondrial reactive oxygen species. Mol Cell. 2012; 48(2):158–67. doi: 10.1016/j.molcel.2012.09.025

38. Yamamoto M, Kim M, Imai H, Itakura Y, Ohtsuki G. Microglia-Triggered Plasticity of Intrinsic Excitability Modulates Psychomotor Behaviors in Acute Cerebellar Inflammation. Cell Rep. 2019; 28(11):2923–2938.e8. doi: 10.1016/j.celrep.2019.07.078

39. Yamamoto Sakai M, Yu Z, Taniguchi M, Picotin R, Oyama N, Stellwagen D, Ono C, Kikuchi Y, Matsui K, Nakanishi M, Yoshii H, Furuyashiki T, Abe T, Tomita H. N-Acetylcysteine Suppresses Microglial Inflammation and Induces Mortality Dose-Dependently via Tumor Necrosis Factor-α Signaling. Int J Mol Sci. 2023;24(4):3798. doi: 10.3390/ijms24043798

40. Janova H, Böttcher C, Holtman IR, Regen T, van Rossum D, Götz A, Ernst AS, Fritsche C, Gertig U, Saiepour N, Gronke K, Wrzos C, Ribes S, Rolfes S, Weinstein J, Ehrenreich H, Pukrop T, Kopatz J, Stadelmann C, Salinas-Riester G, Weber MS, Prinz M, Brück W, Eggen BJ, Boddeke HW, Priller J, Hanisch UK. CD14 is a key organizer of microglial responses to CNS infection and injury. Glia. 2016; 64(4):635–49. doi: 10.1002/glia.22955

41. Beschorner R, Schluesener HJ, Gözalan F, Meyermann R, Schwab JM. Infiltrating CD14+ monocytes and expression of CD14 by activated parenchymal microglia/macrophages contribute to the pool of CD14+ cells in ischemic brain lesions. J Neuroimmunol. 2002 May;126(1-2):107–15. doi: 10.1016/s0165-5728(02)00046-2

42. Ruan C, Elyaman W. A New Understanding of TMEM119 as a Marker of Microglia. Front Cell Neurosci. 2022 Jun 13;16:902372. doi: 10.3389/fncel.2022.902372

43. Simard AR, Soulet D, Gowing G, Julien JP, Rivest S. Bone marrow-derived microglia play a critical role in restricting senile plaque formation in Alzheimer’s disease. Neuron. 2006;49:489–502. doi: 10.1016/j.neuron.2006.01.022.

44. Pangrazzi L, Genovesi S, Balasco L, Cerilli E, Robol C, Zunino G, Piazza S, Provenzano G, Bozzi Y. Immune dysfunction in the cerebellum of mice lacking the autism candidate gene Engrailed 2. J Neuroimmunol. 2022;367:577870. doi: 10.1016/j.jneuroim.2022.577870

45. Love MI, Huber W, Anders S. Moderated estimation of fold change and dispersion for RNA-seq data with DESeq2. Genome Biol. 2014;15(12):550. doi: 10.1186/s13059-014-0550-8

46. Anders S, Huber W. Differential expression analysis for sequence count data. Genome Biol. 2010;11(10):R106. doi: 10.1186/gb-2010-11-10-r106

47. McCarthy DJ, Chen Y, Smyth GK. Differential expression analysis of multifactor RNA-Seq experiments with respect to biological variation. Nucleic Acids Res. 2012;40(10):4288–97. doi: 10.1093/nar/gks042

48. Tenório MCDS, Graciliano NG, Moura FA, Oliveira ACM, Goulart MOF. N-Acetylcysteine (NAC): Impacts on Human Health. Antioxidants (Basel). 2021; 10(6):967. doi: 10.3390/antiox10060967

49. Moy SS, Nadler JJ, Perez A, Barbaro RP, Johns JM, Magnuson TR, Piven J, Crawley JN. Sociability and preference for social novelty in five inbred strains: an approach to assess autistic-like behavior in mice. Genes Brain Behav. 2004; 3(5):287–302. doi: 10.1111/j.1601-1848.2004.00076.x

50. Pastore A, Panera N, Mosca A, et al. Changes in Total Homocysteine and Glutathione Levels After Laparoscopic Sleeve Gastrectomy in Children with Metabolic-Associated Fatty Liver Disease. Obesity Surgery. 2022 Jan;32(1):82–89. DOI: 10.1007/s11695-021-05701-6

