## Supplemental Tables for "The interplay between oxidative stress and inflammation supports autistic-related behaviors in mice"

**Table S1. Summary of RT-PCR data for inflammation and tissue damage related mRNAs in the *Cntnap2^-/-^* cerebellum**

| **Gene name** | **FC**  ***Cntnap2^-/-^***  **vs**  ***Cntnap2^+/+^*** | **p value** |
| --- | --- | --- |
| IL-6 | 1,8 | **** |
| TNF | 1,4 | ** |
| IFNγ | 2,2 | **** |
| IL-1β | 1,5 | **** |
| IFNβ | 2,1 | **** |
| IL-15 | 1,1 | ns |
| TLR1 | 1,4 | ** |
| TLR2 | 2,1 | *** |
| TLR3 | -1,1 | ns |
| TLR4 | 1,1 | ns |
| TLR5 | 2,5 | **** |
| TLR6 | 1,3 | *** |
| TLR9 | 1,9 | **** |
| CCL2 | 1 | ns |
| CCL3 | 2 | **** |
| CCL5 | 2,3 | * |
| CCL8 | 2 | **** |
| CCL21A | 1,9 | **** |
| CCL20 | 1,8 | **** |
| MMP3 | 2,3 | **** |
| MMP8 | 1,4 | *** |
| MMP12 | 2,1 | **** |
| MMP13 | 1,6 | *** |

**Table S2. mRNAs upregulated in the *Cntnap2^-/-^* cerebellum**

| **external_gene_name** | **log2FoldChange** | | **lfcSE** | | **stat** | | **pvalue** | | **padj** |
| --- | --- | --- | --- | --- | --- | --- | --- | --- | --- |
| Slc6a3 | 2,150218872 | | 0,762524 | | 2,81987 | | 0,004804 | | 0,042821 |
| S100a8 | 1,44988535 | | 0,607091 | | 2,388249 | | 0,016929 | | 0,040519 |
| S100a9 | 1,326231844 | | 0,551229 | | 2,405955 | | 0,01613 | | 0,048278 |
| Rpl17-ps5 | 1,030605405 | | 0,310896 | | 3,314955 | | 0,000917 | | 0,017057 |
| Ccr1 | 0,981425315 | | 0,286166 | | 3,429564 | | 0,000605 | | 0,013698 |
| Rpl24 | 0,970737507 | | 0,267496 | | 3,628981 | | 0,000285 | | 0,009409 |
| Pla2g4c | 0,888299819 | | 0,225455 | | 3,940031 | | 8,15E-05 | | 0,00455 |
| Rpl9-ps6 | 0,881334999 | | 0,221831 | | 3,973002 | | 7,1E-05 | | 0,004447 |
| Olfr922 | 0,870362999 | | 0,298647 | | 2,914351 | | 0,003564 | | 0,035985 |
| Il22 | 0,867566172 | | 0,190303 | | 4,558867 | | 5,14E-06 | | 0,001174 |
| Rps6-ps4 | 0,819504243 | | 0,192996 | | 4,246233 | | 2,17E-05 | | 0,002241 |
| Il22b | 0,818251187 | | 0,168531 | | 4,855194 | | 1,20E-06 | | 0,000609 |

**Table S3. mRNAs downregulated in the *Cntnap2^-/-^* cerebellum**

| **external_gene_name** | **log2FoldChange** | **lfcSE** | **stat** | **pvalue** | **padj** |
| --- | --- | --- | --- | --- | --- |
| Cntnap2 | -2,717737225 | 0,144405 | -18,8202 | 5,16E-79 | 8,61E-75 |
| Pax8 | -1,436984411 | 0,490718 | -2,92833 | 0,003408 | 0,034999 |
| Hlx | -1,435347293 | 0,410977 | -3,49253 | 0,000478 | 0,012311 |
| Ddn | -1,408779106 | 0,412051 | -3,41894 | 0,000629 | 0,014016 |
| Kcnj4 | -1,230120725 | 0,30689 | -4,00834 | 6,11E-05 | 0,004084 |
| Cysrt1 | -1,227257755 | 0,287774 | -4,26465 | 2,00E-05 | 0,002143 |
| Nxph2 | -1,182922851 | 0,459606 | -2,57378 | 0,01006 | 0,037006 |
| Scand1 | -1,16902485 | 0,360682 | -3,24115 | 0,00119 | 0,019615 |
| Slc39a4 | -1,139552402 | 0,350705 | -3,24932 | 0,001157 | 0,019316 |
| Tbr1 | -1,052206627 | 0,347648 | -3,02665 | 0,002473 | 0,029538 |
| Mafa | -1,018815166 | 0,341461 | -2,9837 | 0,002848 | 0,032061 |
| Egr2 | -1,009749617 | 0,332221 | -3,03939 | 0,002371 | 0,028771 |

**Table S4. Summary of RNAseq data for inflammation, oxidative stress,**

**and complement mRNAs in the *Cntnap2^-/-^* cerebellum**

| **external_gene_name** | **log2FoldChange** | **lfcSE** | **stat** | **pvalue** | **padj** |
| --- | --- | --- | --- | --- | --- |
| Ccr1 | 0,981425315 | 0,286166 | 3,429564 | 0,000605 | 0,013698 |
| Il22 | 0,867566172 | 0,190303 | 4,558867 | 5,14E-06 | 0,001174 |
| Il22b | 0,818251187 | 0,168531 | 4,855194 | 1,20E-06 | 0,000609 |
| Pla2g2f | 0,698821189 | 0,250654 | 2,787986 | 0,005304 | 0,045372 |
| Ctsw | 0,698461902 | 0,222914 | 3,133319 | 0,001728 | 0,024234 |
| Ccl27a | 0,487455432 | 0,162707 | 2,995907 | 0,002736 | 0,031383 |
| Cat | -0,184164799 | 0,057816 | -3,18537 | 0,001446 | 0,021961 |
| Sod3 | -0,457528115 | 0,126116 | -3,62783 | 0,000286 | 0,009432 |
| Mmp15 | -0,519294807 | 0,154911 | -3,35222 | 0,000802 | 0,016187 |
| C4b | -0,584036148 | 0,147311 | -3,96466 | 7,35E-05 | 0,004447 |
| C1qtnf1 | -0,687953712 | 0,169037 | -4,06985 | 4,7E-05 | 0,003534 |
| Itgax | -0,800650571 | 0,256999 | -3,11539 | 0,001837 | 0,02488 |
| C3 | -0,813910271 | 0,247288 | -3,29135 | 0,000997 | 0,017732 |

|  | **Table S5. Effect of NAC treatment on inflammation/oxidative stress/tissue damage mRNAs in the cerebellum of *Cntnap2^+/+^* and *Cntnap2^-/-^* mice (RT-qPCR data).** | | | | | | |
| --- | --- | --- | --- | --- | --- | --- | --- |
| **Gene** | | **FC**  ***Cntnap2^-/-^* PBS**  ***vs***  ***Cntnap2^+/+^* PBS** | **FC**  ***Cntnap2^-/-^* NAC *vs***  ***Cntnap2^+/+^PBS*** | | | **FC**  ***Cntnap2^-/-^*NAC**  ***vs***  ***Cntnap2^-/-^*PBS** | **FC**  ***Cntnap2^+/+-^*NAC**  ***vs***  ***Cntnap2^+/+^*PBS** |
| **CCL2** | | 4.79 | | -1.95 | -9.38 | | 1.02 |
| **IL-6** | | 18.2 | | 1.13 | -16.1 | | 3.31 |
| **IL-1β** | | 1.93 | | -1.08 | -2.12 | | -1.09 |
| **CCL8** | | -1.39 | | -5.74 | -3.24 | | -3.23 |
| **CCL3** | | 11.7 | | -1.08 | -12.1 | | 2.06 |
| **CCL5** | | 6.69 | | -1.54 | -10.3 | | 1.72 |
| **TNF** | | 28.1 | | 1.19 | -23.5 | | 4.52 |
| **IFNγ** | | 40.2 | | 1.41 | -27.2 | | 6.16 |
| **IFN-β** | | 60.0 | | 1.10 | -29.1 | | 8.54 |
| **CCL21A** | | 5.03 | | 1.33 | -3.77 | | 1.18 |
| **CCL20** | | 58.7 | | 2.35 | -24.9 | | 7.93 |
| **TLR1** | | 12.8 | | 1.21 | -10.6 | | 2.48 |
| **TLR2** | | 1.06 | | -1.24 | -1.31 | | -1.17 |
| **TLR3** | | 1.09 | | -1.08 | -1.18 | | -1.16 |
| **TLR4** | | 3.19 | | 1.14 | -2.81 | | 10.9 |
| **TLR5** | | 17.0 | | 1.25 | -13.6 | | 2.64 |
| **TLR9** | | 7.18 | | 1.09 | -6.56 | | 1.57 |
| **MMP3** | | 35.9 | | 1.44 | -24.8 | | 5.07 |
| **MMP8** | | 1.44 | | 1.52 | 1.07 | | 1.07 |
| **MMP12** | | 18.7 | | -1.11 | -20.7 | | 2.89 |
| **MMP13** | | 1.37 | | -1.51 | -1.97 | | -1.28 |
| **IL-22** | | 1.47 | | 1.15 | -1.22 | | 1.04 |
| **CCL12** | | 26.4 | | 2.05 | -12.9 | | 3.64 |
| **CCL27A** | | 1.57 | | 1.21 | -1.29 | | -1.04 |
| **SOD3** | | -1.74 | | -2.82 | -1.62 | | -2.29 |

**Table S6. Sequences of primers used for RT-qPCR experiments.**

| Target gene | Forward primer (5’-3’) | Reverse primer (5’-3’) |
| --- | --- | --- |
| IL-6 | GCCTTCTTGGGACTGATGCT | GACAGGTCTGTTGGGAGTGG |
| IL-1β | ACGGACCCCAAAAGATGAAG | TTCTCCACAGCCACAATGAG |
| TNF | CAAAATTCGAGTGACAAGCC | TGTCTTTGAGATCCATGCCG |
| IFNγ | CCCTATGGAGATGACGGAGA | CTGTCTGCTGGTGGAGTTCA |
| IFNβ | CGAGCAGAGATCTTCAGGAAC | TCACTACCAGTCCCAGAGTC |
| IL-15 | ATCCATCTCGTGCTACTTGTGTT | CATCTATCCAGTTGGCCTCTGTTT |
| CCL2 | GAGTAGGCTGGAGAGCTACAAGAG | AGGTAGTGGATGCATTAGCTTCAG |
| CCL3 | TGAAACCAGCAGCCTTTGCT | AGGCATTCAGTTCCAGGTCAGTG |
| CCL5 | AGAATACATCAACTATTTGGAGA | CCTTGCATCTGAAATTTTAATGA |
| CCL8 | GCTGTGGTTTTCCAGACCAA | GAAGGTTCAAGGCTGCAGAA |
| CCL20 | CTTGCTTTGGCATGGGTACT | TCAGCGCACACAGATTTTCT |
| CCL21A | ATGGCTCAGATGATGACTCTGAGC | GTACTTAAGGCAGCAGTCCTGA |
| MMP3 | CAGACTTGTCCCGTTTCCAT | GGTGCTGACTGCATCAAAGA |
| MMP12 | GGAGCTCACGGAGACTTCAACT | CCTTGAATACCAGGTCCAGGATA |
| MMP13 | CTTGATGCCATTACCAGTC | GGTTGGGAAGTTCTGGCCA |
| TLR1 | TCAAGTGTGCAGCTGATTGC | TAGTGCTGACGGACACATCC |
| TLR2 | TGATGGTGAAGGTTGGACG | CGGAGGGAATAGAGGTGAAAG |
| TLR3 | CCTCCAACTGTCTACCAGTTCC | GCCTGGCTAAGTTATTGTGC |
| TLR4 | CAACATCATCCAGGAAGGC | GAAGGCGATACAATTCCACC |
| TLR5 | AGCATTCTCATCGTGGTGG | AATGGTTGCTATGGTTCGC |
| TLR6 | TGGATGTCTCACACAATCGG | GCAGCTTAGATGCAAGTGAGC |
| TLR9 | CAAGAACCTGGTGTCACTGC | TGCGATTGTCTGACAAGTCC |
| IL-22 | CGATTGGGGAACTGGACCTG | GGACGTTAGCTTCTCACTTT |
| CCL12 | ATCCAGAGCTTGAGTGTGACGC | AAGGCAAACTTTTTGACCGCC |
| CCL27A | TTCCTTGGCTGCGAATGT | CTTCTGCTTAGTCTTGTTCCA |
| SOD3 | CCAGCTTCGACCTAGCAGACA | CAGCGTGGCTGATGGTTGTA |
| TMEM119 | GTGTCTAACAGGCCCCAGAA | AGCCACGTGGTATCAAGGAG |
| β actin | GGCTGTATTCCCCTCCATCG | CCAGTTGGTAACAATGCCATGT |

**Table S7. Antibodies used for the flow cytometry experiments**

| Antigen | Fluorocrome | Company | Clone |
| --- | --- | --- | --- |
| CD45 | PE | Miltenyi | REA747 |
| CD44 | FITC | Biolegend | IM7 |
| CD3 | APC-Vio770 | Miltenyi | REA641 |
| CD8 | PerCp | Biolegend | 53–6.7 |
| CD62L | APC | Biolegend | MEL-14 |
| CD4 | VioGreen | Miltenyi | REA1211 |
| CD14 | Pecy7 | Biolegend | Sa14–2 |
| IL-6 | APC | Biolegend | MP5-20F3 |
| IFNγ | FITC | Miltenyi | REA638 |
| TNF | PE | Biolegend | MP6-XT22 |
| IL-2 | APC | Biolegend | JES6-5H4 |
| Tmem119 | Pecy7 | ThermoF. | V3RT1GOsz |
| Neun | FITC | Miltenyi | REA1131 |
