## Supplementary Figures for "The interplay between oxidative stress and inflammation supports autistic-related behaviors in mice"

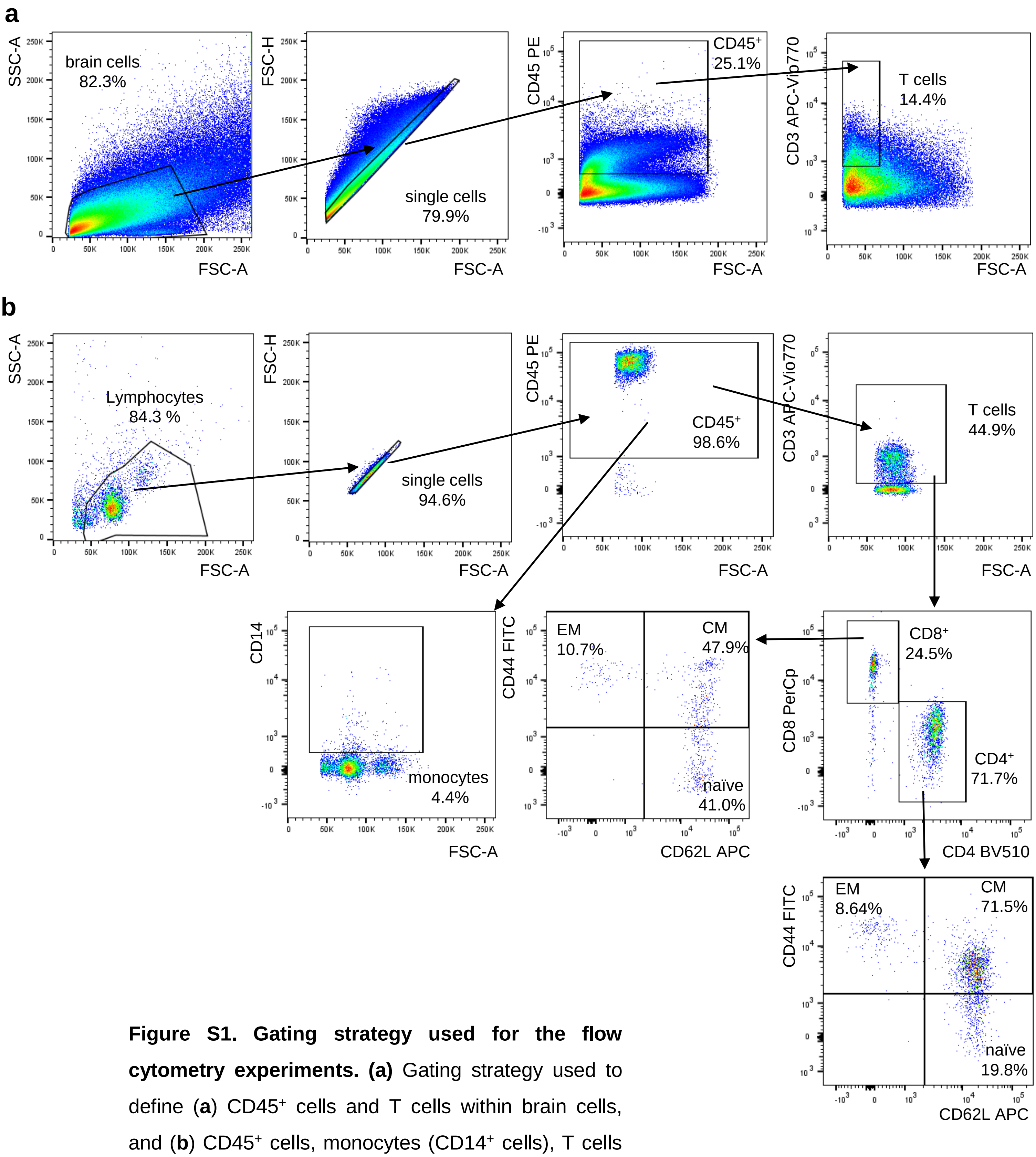


**Figure S1. Gating strategy used for the flow cytometry experiments. (a)** Gating strategy used to define (**a**) CD45⁺ cells and T cells within brain cells, and (**b**) CD45⁺ cells, monocytes (CD14⁺ cells), T cells (CD8⁺ and CD4⁺) and CD44⁺CD62L⁻ EM, CD44⁺CD62L⁺ CM and CD44⁻CD62L⁺ naïve cells in CD8⁺ and CD4⁺ T cells within PBMCs.

### Supplementary Figure 1

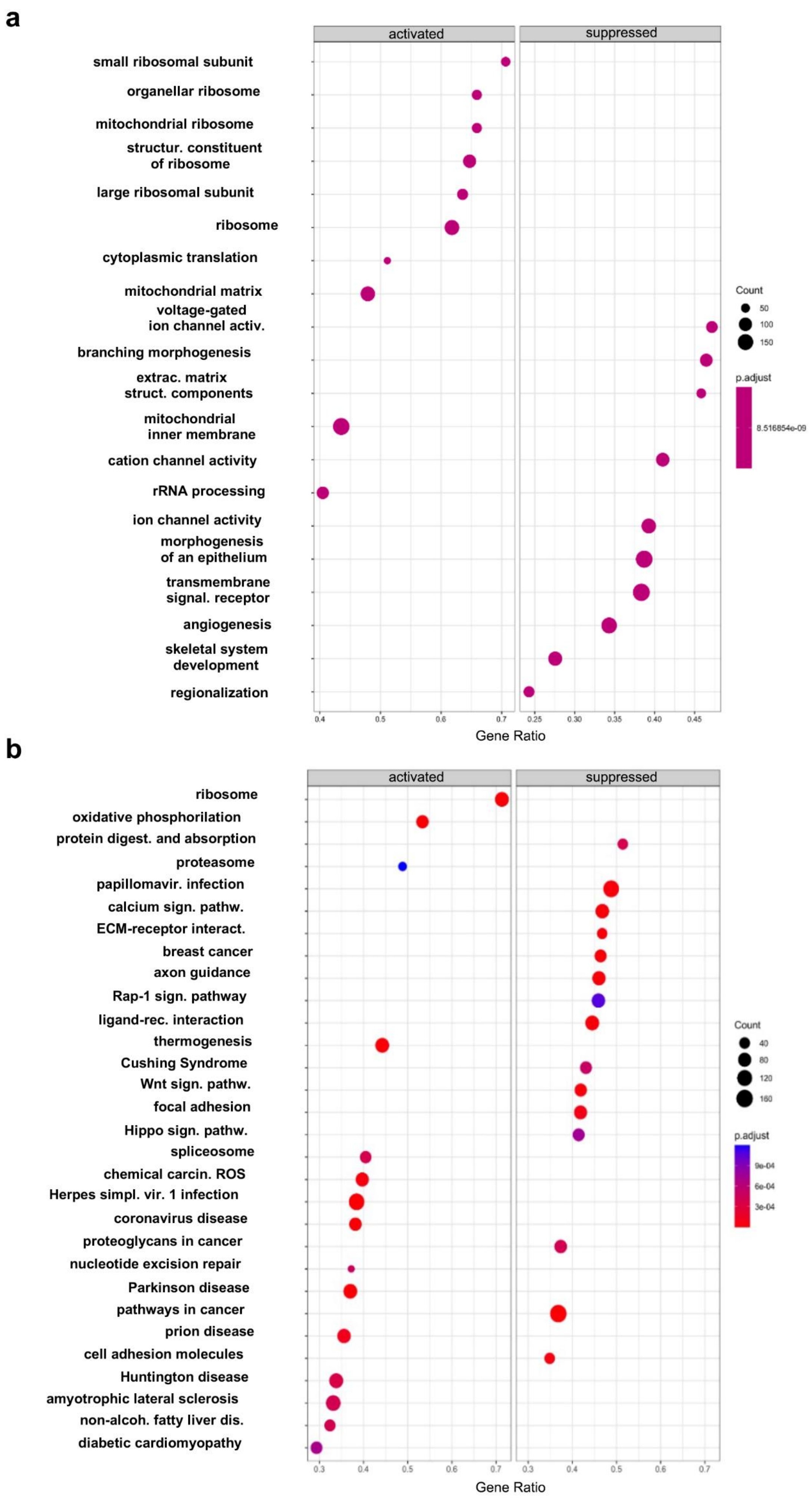


### Supplementary Figure 2

**Figure S2. GO and KEGG pathway analysis of RNAseq data.** Positively and negatively enriched (a) GO and (b) KEGG pathways obtained after the GSEA.

**a**

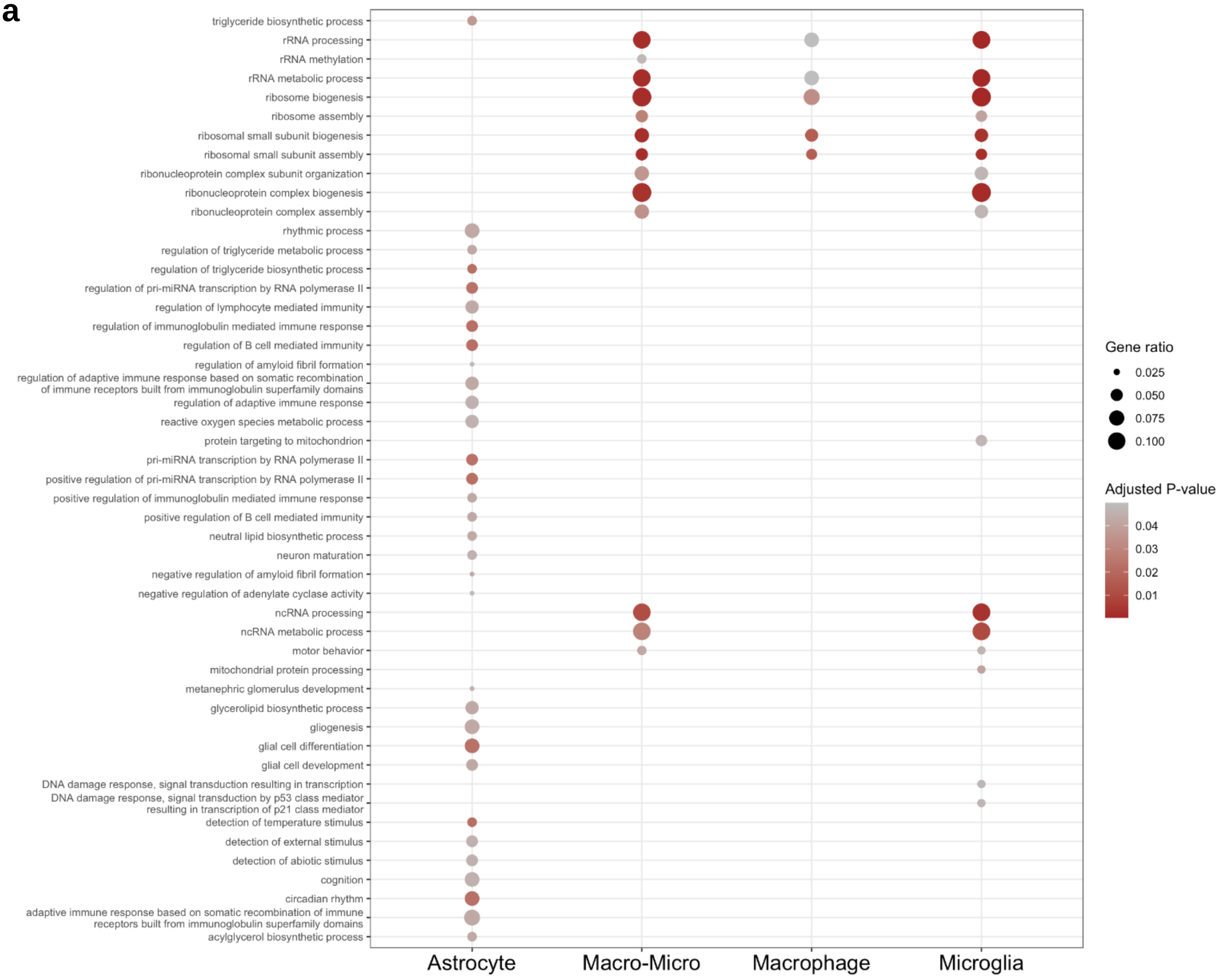


**b**

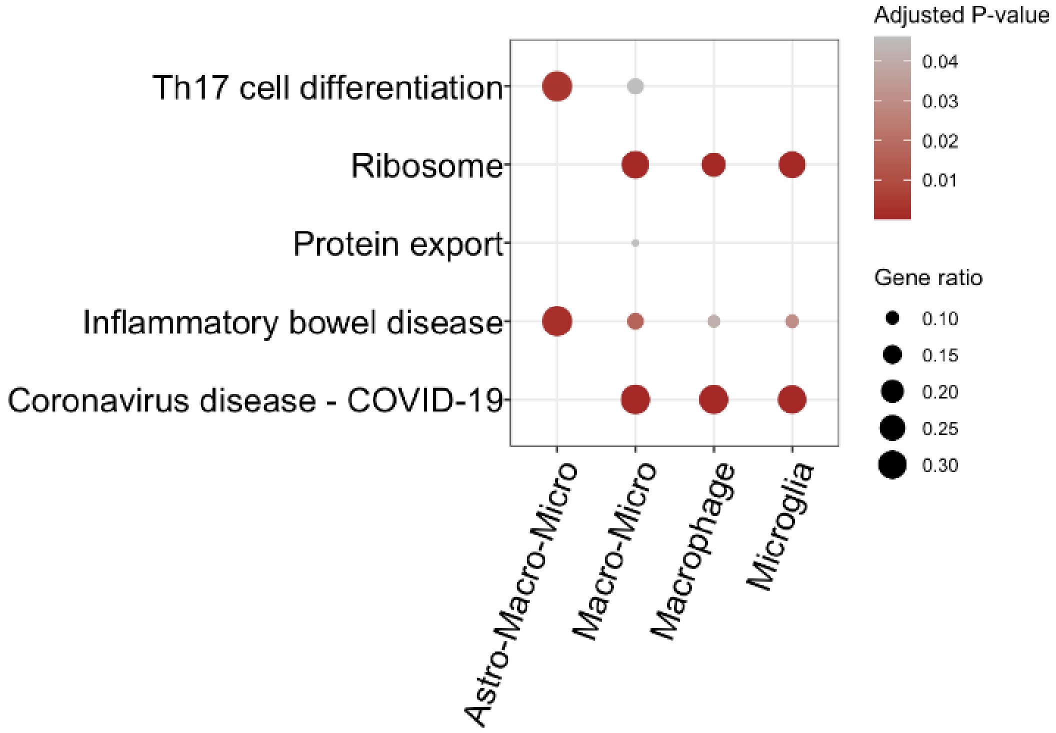


### Supplementary Figure 3

**Figure S3. GO and KEGG pathway analysis of RNAseq data after cell deconvolution analysis.** (**a**) GO and (**b**) KEGG pathway analysis after adjusting for cell subsets (astrocytes, macrophages+microglia, macrophages, and microglia. Adjusted p values are displayed in the graphs using the scale color.

### Supplementary Figure 4

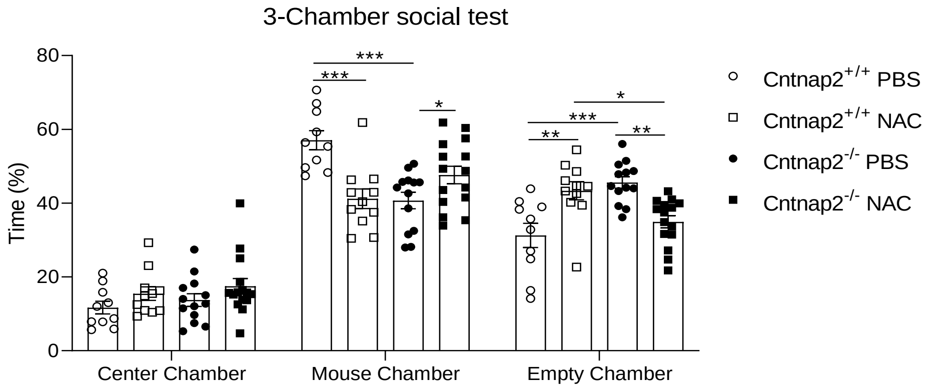


**Figure S4. Percentage of time in the 3 chambers in the 3-chamber test.** Percentage of time in the center chamber, mouse chamber and empty chamber in the 3-chamber social test. Two-way ANOVA, Tukey post-hoc test. *p<0.05; **p<0.01; ***p<0.001. $n_{Cntnap2+/+PBS=}7$, $n_{Cntnap2+/+NAC=}6$, $n_{Cntnap2-/-PBS=}7$, $n_{Cntnap2+/+NAC=}8$. Two-way ANOVA, Tukey post-hoc test. *p<0.05; ****p<0.0001.

### Supplementary Figure 5

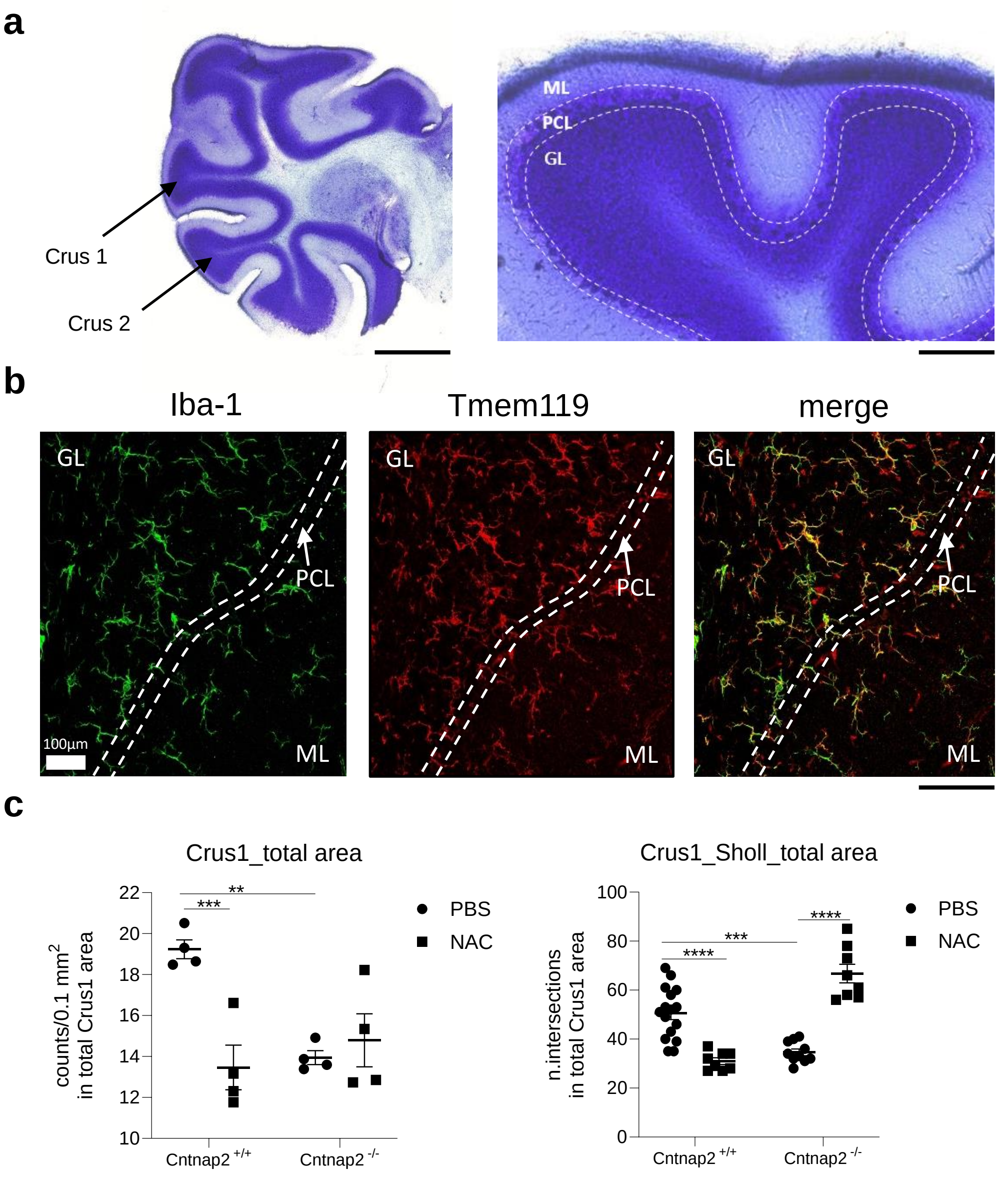


**Figure S5. Analysis of microglia cells in the cerebellum of PBS- and NAC- treated mice.** (**a**) representative picture of Nissl staining in the cerebellum of *Cntnap2*$^{+/+}$ mice. Crus1 and Crus 2 areas, with their respective molecular layer (ML), Purkinje cells layer (PCL) and granular layer (GL) are displayed. (**b**) Representative images showing the colocalization between iba-1$^{+}$ and tmem119$^{+}$ cells within Crus2 area. (**c**) iba-1$^{+}$ cells (expressed in 0.1 mm2) and (**d**) intersections calculated with the Sholl analysis in the Crus1 total area. For cell number quantification: n=4 in each group; For Sholl analysis: $n_{Cntnap2+/+PBS=}$16, $n_{Cntnap2+/+NAC=}$8, $n_{Cntnap2-/-PBS=}$10, $n_{Cntnap2+/+NAC=}$8. Two-way ANOVA, Tukey post-hoc test. *p<0.05; **p<0.01; ***p<0.001; ****p<0.0001.

### Supplementary Figure 6

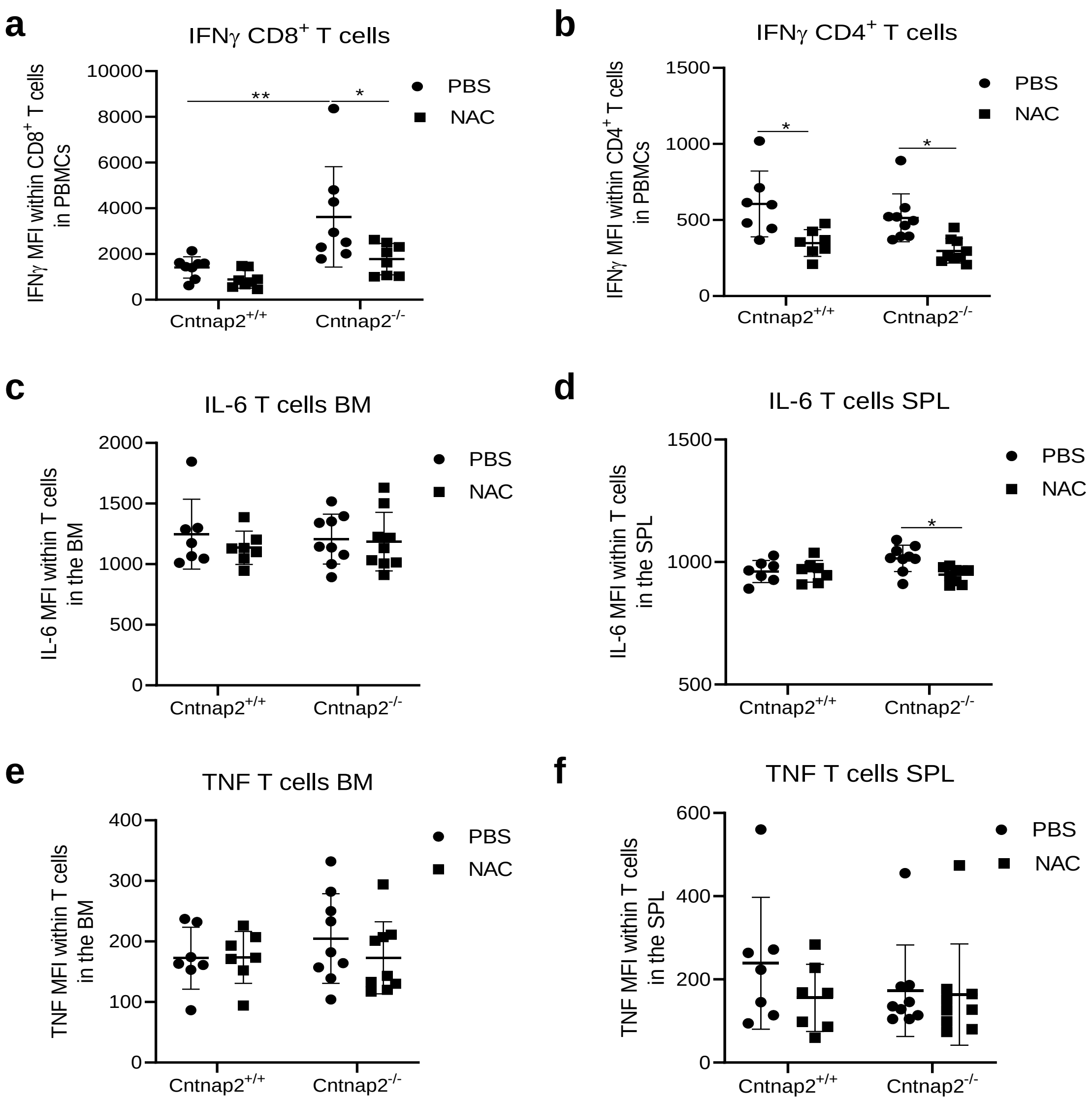


**Figure S6. The impact of NAC on pro-inflammatory molecules within T cell subsets in the peripheral blood, bone marrow, and spleen of PBS- and NAC-treated *Cntnap2$^{+/+}$* and *Cntnap2$^{-/-}$* mice.** Mean fluorescence intensity (MFI) of (a) IFNγ in $CD8^+$ T cells, and (b) IFNγ in $CD4^+$ T cells within PBMCs. MFI of IL-6 in $CD8^+$ T cells within (c) the bone marrow, and (d) the spleen. MFI of TNF in $CD8^+$ T cells within (e) the bone marrow, and (f) the spleen. n=7-9 in each group. Two-way ANOVA, Tukey post-hoc test. *$p<0.05$; **$p<0.01$.
